# The genomic landscape of wild *Saccharomyces cerevisiae* is shaped by complex patterns of admixture, aneuploidy and recombination

**DOI:** 10.1101/2023.06.07.544145

**Authors:** Chris M. Ward, Cristobal A. Onetto, Steven Van Den Heuvel, Kathleen M. Cuijvers, Laura J. Hale, Anthony R. Borneman

## Abstract

Cultural exchange of fermentation techniques has driven the spread of *Saccharomyces cerevisiae* across the globe, establishing wild populations in many countries. Despite this, most modern commercial fermentations are inoculated using monocultures, rather than relying on natural populations, potentially impacting wild population diversity. Here we investigate the genomic landscape of 411 wild *S. cerevisiae* isolated from spontaneous grape fermentations in Australia across multiple locations, years, and grape cultivars. Spontaneous fermentations contained highly recombined mosaic strains that commonly exhibited aneuploidy of chromosomes 1, 3, 6 and 9. Assigning wild genomic windows to putative ancestral origin revealed that few closely related commercial lineages have come to dominate the genetic landscape, contributing most of the genetic variation. Fine-scale phylogenetic analysis of loci not observed in strains of commercial wine origin identified widespread admixture with the Beer2 clade along with three independent admixture events from potentially endemic Oceanic lineages that last shared an ancestor with modern East Asian *S. cerevisiae* populations. Our results illustrate how commercial use of microbes can affect local microorganism genetic diversity and demonstrates the presence of non-domesticated, non-European derived lineages of *S. cerevisiae* in Australian ecological niches that are actively admixing.

## Introduction

The yeast *Saccharomyces cerevisiae* has been tied to human culture and movement since Neolithic communities in the Middle- and Far-East began fermenting fruits and grain 8000-9000 years ago^1,2^. Phylogenomic evidence further supports this, with natural isolates from forests of modern China and South East Asia forming the crown node to all domesticated isolates of *S. cerevisiae*^*3,4*.^ Grapes and their associated fermentation products are then thought to have spread through trade and dispersion into the Near East and East Mediterranean regions^5^, resulting in the endemic wild populations identified in China (CN)^3^; Taiwan (TW)^6^; Japan^7^; Europe^7^; North America (NA)^7^; Africa^8^ and South East Asia^8^ forming clear taxonomic clades^3,4,6,8^.

Historically, wine was fermented spontaneously, with inoculation by autochthonous (native/endemic/wild) yeast occurring through environmental transfer^9-11^. This had the potential to confer unique and terroir-dependent sensory traits to wine that would otherwise be absent^12-14^. In contrast, most modern industrial-scale fermentations are inoculated using commercial starter cultures containing a single yeast monoculture^9,15,16^, providing predictability of fermentation characteristics^12,17^; we speculated that frequency and quantity of commercial inoculations may have significant effects on the wild genetic landscape and strain biodiversity.

Admixture and horizontal gene transfer has been essential in driving the diversification of S. cerevisiae^18-20^, with admixture from basal Asian isolates into wine populations being proposed as likely the origin of the Beer2 clade that is associated with modern ale production^18^. Widespread use, and subsequent dispersal, of domesticated strains has been a major contributor to the frequency of admixture^21^ and has led to the establishment of diverse wild populations throughout the world. Wild *S. cerevisiae* have been isolated in low frequencies from the guts of insects, grape vines, and fermentation related machinery, providing multiple distinct avenues for wild isolates to be inoculated into the fermentation medium^7,20,22,23^. Despite this, key questions regarding the genetic composition of wild *S. cerevisiae* populations remain.

In this study, whole-genome resequencing was performed on 411 *S. cerevisiae* strains that were isolated from spontaneous fermentations from grape varieties across a five-year period in Australian wineries. Through comparison against 169 commercial wine isolates and 91 diverse yeast isolates^3,7,8^, the genomic landscape of wild yeast shows evidence of frequent aneuploidy, recombination between commercial haplotypes and admixture events from basal potentially endemic lineages. Furthermore, after screening cultures of Australian wild niches a single isolate was identified from that was monophyletic with introgressed loci and the NA & Japan clade.

## Results

### Isolate origin and genotyping

Extracts from spontaneous fermentations from five grape varieties (Viognier, Pinot Noir, Shiraz, Grenache, Chardonnay and Cabernet Sauvignon) were cultured and isolates with ITS sequences matching *Saccharomyces cerevisiae* (n = 434) were sequenced using short read Illumina platforms outputting an estimated mean coverage of 29.9x (95% CI: 28.5x-31.3x) per isolate. Short reads were investigated for species origin using exact 31-mer matches against the genomes of *sensu stricto Saccharomyces* and out of 434 sequenced wild isolates, 427 were classified as *S. cerevisiae* (Sup Figure 1). Furthermore, no significant signal for between species hybridisation was identified, with >99% of classified samples k-mers matching *S. cerevisiae*. After filtering for average read depth (DP≥6) and genotyped coverage (>80% of S288C genome^24^), a total of 411 wild isolates, along with 169 commercial wine^16^ and 91 diverse^3,7,8^ publicly available yeast isolates were retained for analysis (Sup Figure 2, Sup Table 1).

### Wild yeast exhibit high levels aneuploidy

Whole genome duplications and aneuploidies have been shown to occur in yeast in response to biotic or abiotic stress and are a hallmark of adaptation to human-associated environments, such as wine and beer production^25,26^. To investigate this, genome-wide ploidy was estimated using frequency bias of biallelic 21-mers for each wild and commercial isolate utilized in this study. This revealed that the majority of both wild and commercial yeast were diploid, while polyploidy occurred at similar frequencies in both wild isolates (8.3%) and their commercial (7.5%) counterparts (Sup Figure 3, Sup Table 2).

Chromosomal aneuploidies were then determined by calculating read depth across 1386 BUSCO genes grouped according to nuclear chromosomes. This produced a dataset containing 2,247,385 sites within single copy orthologs across the S288C genome (29,926-277,678 per chromosome) for analysis. Chromosomes displaying aneuploidy were predicted if the chromosomal depth distribution had a ≥0.5 rank biserial effect size when compared to all other chromosomes based on the results of a Mann-Whitney U-test. Putative events were manually inspected to confirm that all putative events had a read depth bias indicative of whole-chromosome aneuploidy (Figure 1A, Sup Figure 4).

**Figure 1:**
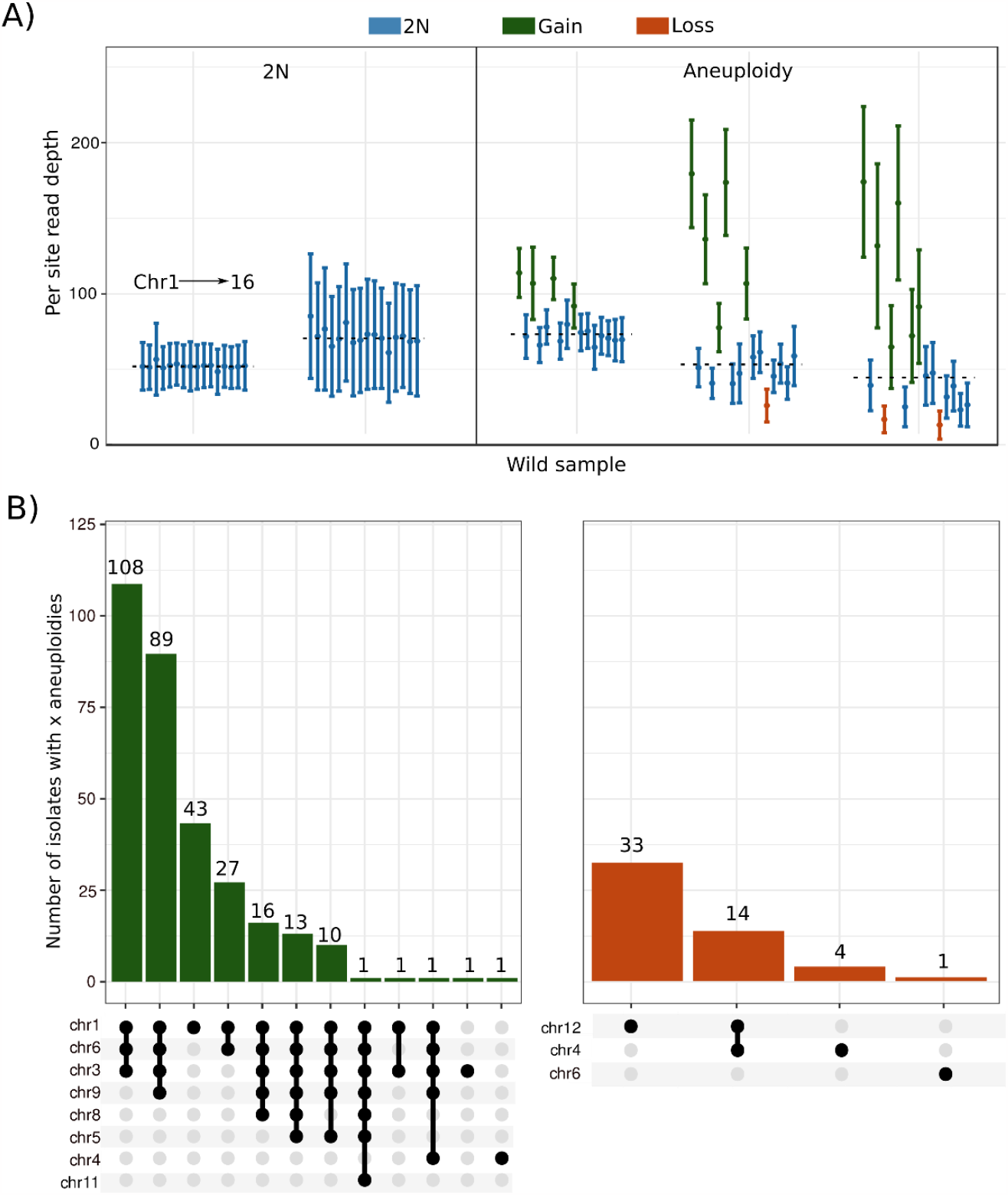
Aneuploidies are widespread in wild yeast. A) Examples of per site read depth within single copy orthologs distributed across chromosomes for two strictly diploid (2N) and three samples displaying combinations of chromosomal aneuploidies. Each point and error bar represents the mean and standard deviation across a *S. cerevisiae* chromosome (1-16). Dashed lines represent the median genome wide depth. Density distributions of within chromosome read depth can be found in Sup Figure 4. B) Chromosome gain (left) and chromosome loss (right) events observed in wild isolates. Upset plot nodes represent aneuploidies that were present simultaneously in the same individual along with the number of times each pattern was observed shown above bars.

Wild isolates displayed a high frequency of aneuploidy, with 75.9% of all strains (Figure 1B, Sup Table 3) containing at least one chromosomal aneuploidy, compared to only 11.2% of commercial strains (Sup Figure 5, Sup Table 4). Chromosomal gain events (Figure 1B; left) were far more frequent than loss (Figure 1B; right), comprising 85.7% of all aneuploidies. A clear negative correlation was also observed between chromosome length and the frequency of chromosomal gain (Sup Figure 6). Most gain events involved chromosomes 1, 3, 6, and 9; with a trend towards multiple aneuploidies being observed within single isolates rather than single aneuploidies being equally distributed through the dataset (Figure 1). Aneuploid chromosomes in wild strains also had a read depth bias indicating higher than 3N copy number in most situations (Figure 1A). In contrast, chromosomal loss events were infrequent and almost exclusively involved the longer chromosomes (Figure 1B), aligning with previous results from natural populations^8^ and artificial selection experiments^26,27^.

### Genetic diversity within wild yeast

Higher heterozygosity rates were observed within individual commercial isolates compared to the wild isolates in this study (Figure 2A), with many wild strains showing extremely low heterozygosity, which may be due to genome renewal^28^. Despite this, average Nei’s nucleotide diversity (*π*)^29^ across the set of wild isolates was greater than that observed across the commercial strains (0.0012±0.0016) in three years (Figure 2B), 2018: 0.0016±0.0024; 2017: 0.0014±0.0021; 2016: 0.0015±0.0025). Although not significant, this suggests that *de novo* mutation; un-sequenced commercial strains; other *S. cerevisiae* clade isolates or other species may be contributing to the standing variation of wild yeast isolates. Furthermore, nucleotide diversity fluctuated between locations (Figure 2B), which was more pronounced when stratifying by location and grape variety, with some locations displaying twice the nucleotide diversity across isolates, even when comparing the same grape variety (Sup Figure 7).

**Figure 2:**
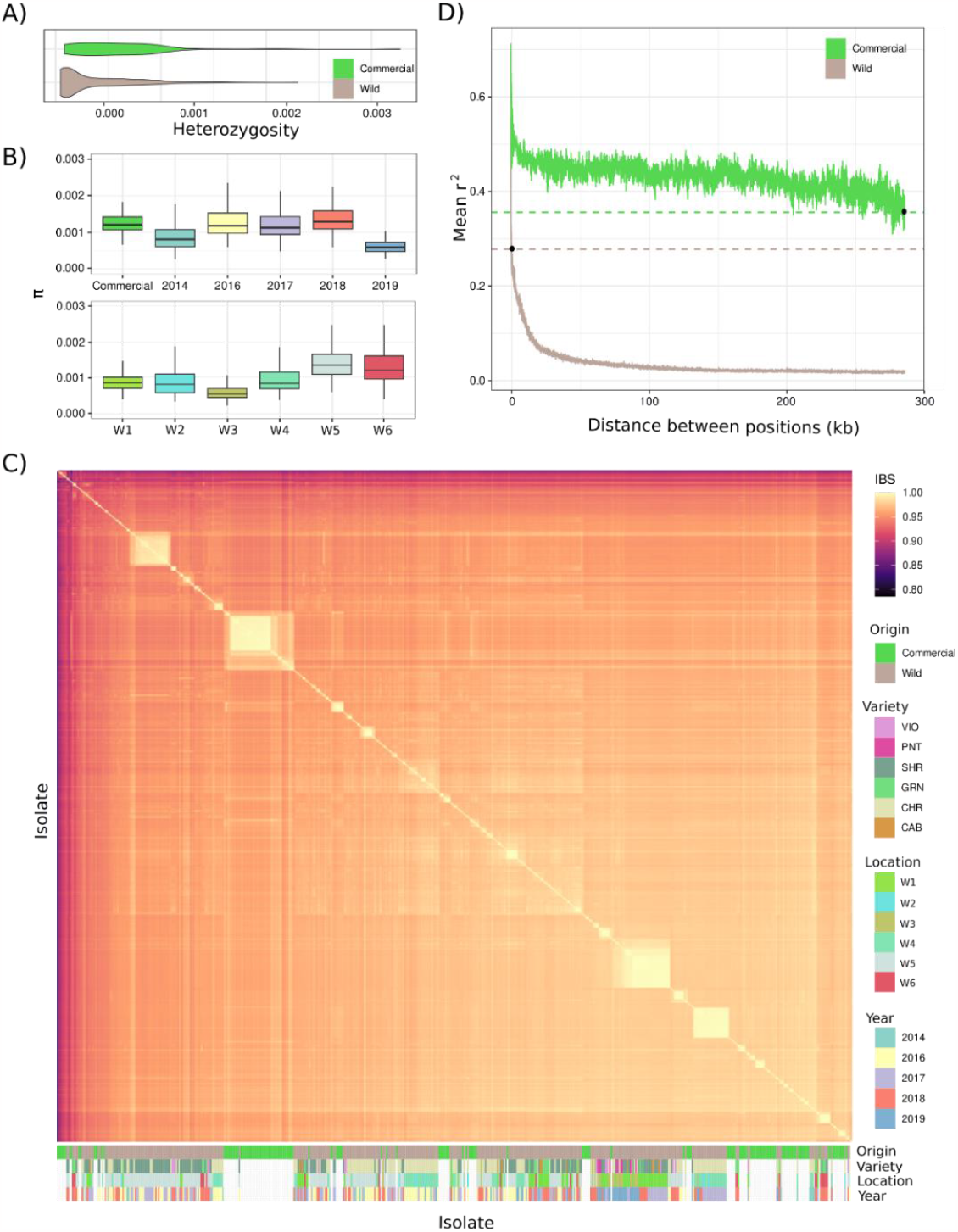
Genetic diversity of commercial and wild yeast isolates. A) Distribution of per isolate genome wide heterozygosity for commercial and wild samples. B) UPPER: Quartiles of 10kb windowed nucleotide diversity (*π*) for commercial or wild yeast collected in 2014, 2016, 2017, 2018 and 2019. LOWER: Quartiles of 10kb windowed nucleotide diversity (*π*) for wild yeast collected from locations W(Winery)1-W6. Black horizontal lines show the median value for the dataset. Boxes showing the first and third quartile range (IQR) while whiskers extend to a maximum of 1.5 * IQR. C) Heatmap of isolate vs isolate identity by descent (IBD) for all Wild and Commercial isolates annotated based on type, variety, location, and year. VIO = Viognier, PNT = Pinot Noir, SHR = Shiraz, GRN = Grenache, CHR = Chardonnay, CAB = Cabernet Sauvignon. D) Decay of linkage disequilibrium for commercial (green) and wild (brown) populations as a function of r_2_ values between pairs of SNPs across the genome. Distances where linkage disequilibrium decayed to half its total value are marked by a black point.

### Recombination between commercial lineages drives wild genome diversity

The simplest explanation for the presence of wild yeast strains is that they represent “escaped” commercial starter strains that have undergone feralization. To determine if this was the case, genome wide identity by state (IBS) calculations were performed between all pairwise comparisons of commercial and wild yeast isolates. While a small subset of wild isolates (n=28) were genetically identical to a specific commercial strain, most were distinct (Figure 2C). Clonal wild haplotype clusters were observed across multiple years and between locations (Figure 2C), suggesting a general lack of population structure within wild yeast that is supported by principal component analysis (PCA) (Sup Figure 8). Interestingly, 33 wild samples isolated from the same location showed low (<0.90) IBS with wine yeast and separated from other wild samples across EV2 (Sup Figure 8). This may be the result of inter-clade hybridization or mixture from divergent wine strains that are not captured in the commercial database used here^16^.

While the wild genotypes did not represent clonal lineages of commercial strains, they could represent recombined genotypes from within the commercial genetic pool. Estimates of linkage disequilibrium (LD) decay between the wild and commercial sets were used to determine if significant sexual recombination was occurring within the wild population. The rate of LD decay was divergent between wild and commercial isolates, with LD decaying to half its maximum value within 0.89 kb and 299.69 kb respectively (Figure 2D). Commercial isolates therefore appear stable, with little sexual recombination, while the wild populations display evidence for extensive recombination.

To determine if the extensive LD decay in wild strains was due to recombination primarily between a set of commercial parental strains, local haplotype origin was investigated across a set of near homozygous wild isolates (genome wide heterozygosity ≤ 0.05%, n = 292). First, hierarchical clustering of the pairwise co-ancestry coefficient (f_(AB)_) between commercial strains was used to reduce the commercial dataset into 25 lineages and 71 unique commercial isolates (Sup Figure 9, Sup Table 5). IBS was then calculated across the wild genomes using IBS (≥0.99) to identify the most probable commercial lineage for 10, 15, 20 and 25kb windows across each wild genome (Figure 3A). All window sizes showed a similar ancestry distribution (Sup Figure 10), therefore the largest (25kb) was selected for further analysis. Wild isolates had an average of 322 (95% CI: 317-326) 25kb windows assigned to commercial origin, with only 16 windows having too few sites (>20%) due to repetitive content. Although few wild isolates were near clones of commercial strains (n = 8, Sup Table 6), the vast majority (99%) could be assigned a mixed commercial ancestry, suggesting that gene-flow and meiotic recombination between commercial haplotypes occurs frequently in the wild (Figure 3A, Sup Figure 11). Most wild isolates were comprised of genetic contributions from many commercial strains and 96% of commercial lineages/isolates contributed genetics to at least one window of a wild isolate genome (Figure 3B, Sup Figure 10). However, one commercial strain 1492 (Enoferm CSM, Lallemand) contributed greater than 0.2 of the genome (95% CI: 0.18 - 0.22) to 118 isolates out of the 292 tested (Figure 3B). Furthermore, lineage l16 contributed an average of 0.12 and was closely related to 1492 (f_(AB)_ ∼ 0.94) (Figure 3C, Sup Figure 9) suggesting these two lineages are contributing to the genetic landscape far more than other commercial isolates. Mosaic wild genomes ranged from slightly (≤ 5 lineages) to highly mixed (≥ 40 lineages) and the genome of each wild strain was derived from a median of 42 different commercial lineages/isolates (Figure 3D). Despite this, many loci failed to be assigned to a sampled commercial isolate at the 0.99 IBS cut-off suggesting these may be the result of gene-flow from outside of the commercial pool utilized in this study or other divergent wild isolates.

**Figure 3:**
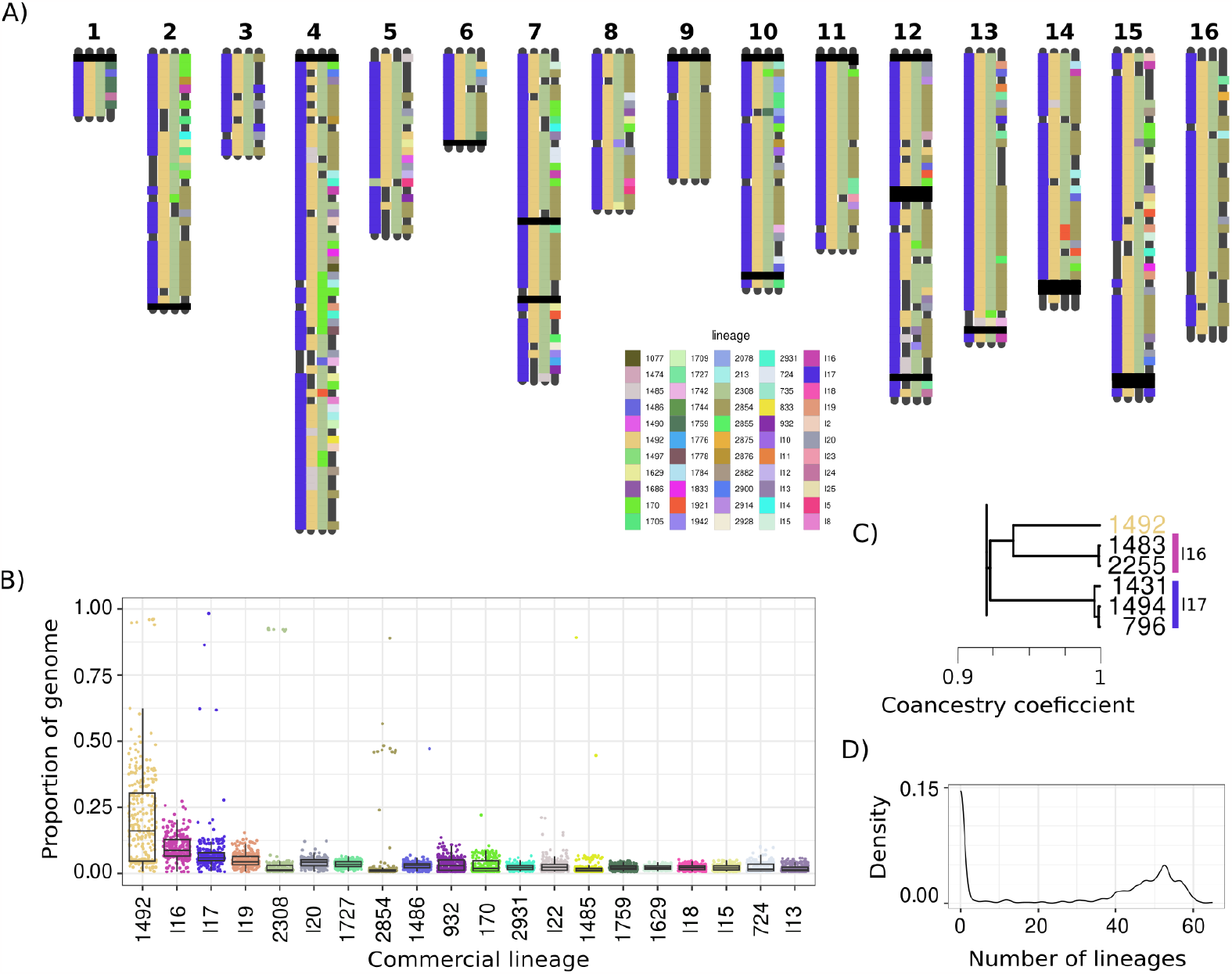
Contribution of commercial strain genetic content to near-homozygous wild genomes. A) Window-wise 25kb Identity by State calculated for all pairwise comparisons of commercial and near-homozygous diploid wild isolates with only select isolates shown. Colors indicate assigned commercial strain lineage of origin ranging from near identical to highly mixed. Block-wise assignment to chromosomes can be found above the figure. An extended figure containing all near-homozygous diploid wild isolates can be found in Sup Figure 11. B) Top 20 commercial lineages that contribute genomic blocks to wild isolates and their proportion contribution to each wild isolate tested based on 25kb windowed IBS. Boxes showing the first and third quartile range (IQR) while whiskers extend to a maximum of 1.5 * IQR. C) Dendrogram showing the co-ancestry relationship between the top three lineages, excised from Sup Figure 9. D) Density plot showing the number of commercial strain lineages that contribute to the genome of lowly heterozygous wild isolates.

### Recent admixture between wild *S. cerevisiae* and non-domesticated isolates

To investigate the genetic provenance of the wild isolates, phylogenetic reconstruction was carried out utilizing the full set of 411 wild and 169 commercial isolates along with a publicly available set of 91 diverse yeast from major domesticated and non-domesticated clades described in Pontes, et al. ^4^. The resultant phylogeny matched previous studies with disparate brewing methods separating into well supported clades (Figure 4A). Commercial wine isolates were mostly monophyletic, bifrucating into the three previously described clades Vin7, PdM and a mixed clade of other wine representatives^16^. However, nine wild isolates were in paraphyly with commercial wine samples suggesting admixture with other ancestral isolates may have occurred. ADMIXTURE analysis with *a priori* population number (K) set from 2-5 was carried out to investigate the drift ancestry of each of the isolates, clearly separating samples into European derived, bounded by the Mediterranean Oak (MO) MRCA, and basal ancestries (Figure 4B, Sup Figure 12). Mixed ancestries were identified for Beer2, MO and Beer1 & Bread clades agreeing with previous work^6,7,18,19^. Paraphyletic and wild isolates at the base of the major European wine clade showed clear mixture of multiple ancestries at all values of K reinforcing the prediction that these are the result of mixed descent (Figure 4B; red boxes).

**Figure 4:**
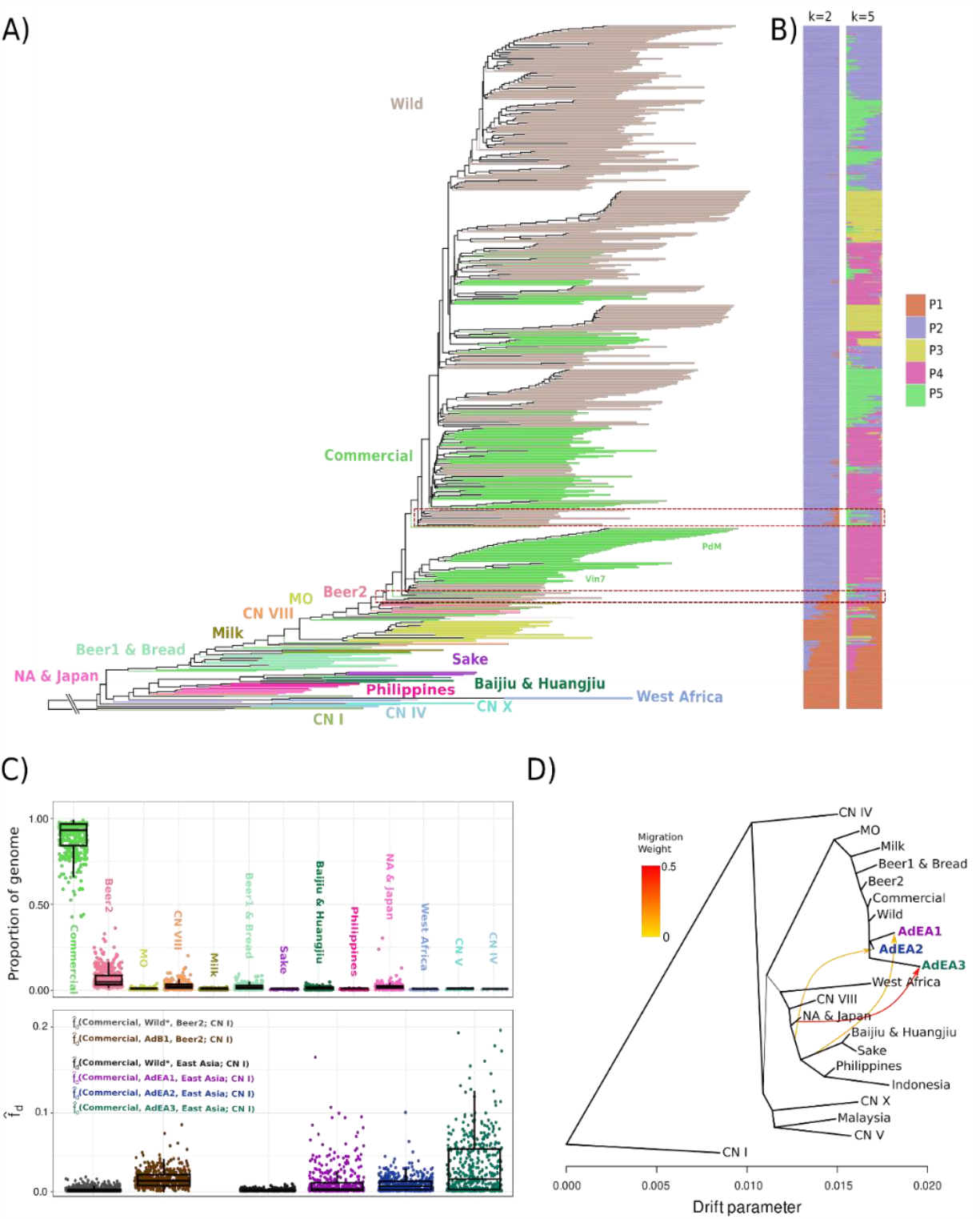
Ancestry inference of wild isolates identified three distinct admixture events from basal Asian clades. A) Whole genome species tree reconstructed from 1386 single copy ortholog gene trees. B) Genome wide estimation of drift histories for k = 2 & 5. C) Upper: The proportion each wild isolates genome that was most closely related to one of the clades in A using 10kb genomic windows of Nei’s d_a_. Lower: *f*_*d*_ tests calculated across 25kb windows to identify allelic bias from Beer2 or East Asia (a combined population of Baijiu & Huangjiu, CN VIII and NA & Japan isolates) present in admixed pseudo-populations that is absent from Commercial isolates. *f*_*d*_ replacing admixed pseudo-populations with Wild* isolates (Wild isolates are not in AdB1, AdEA1, AdEA2 or AdEA3) used as a negative control. Boxes showing the first and third quartile range (IQR) while whiskers extend to a maximum of 1.5 * IQR. D) Drift tree with putative admixed individuals treated as separate populations (AdEA1, AdEA2, AdEA3) allowing for directional ‘migration’ events to simulate admixture between populations. Simplified version of Sup Figure 13 m = 10. All populations are colored based on clade assignment in panel A.

Pairwise Nei’s ancestral distance (*d*_*a*_)^29^ was then calculated across 25kb tiled windows to determine if any wild loci were genetically closer to non-wine isolates than commercial wine isolates (Figure 4C). The proportion of windows most similar to commercial isolates ranged from 0.43 – 0.99 of the total identifiable genetic contribution for each wild isolate (Figure 4C). Non-commercial windows were most associated with the Beer2 clade, contributing between 0.0064 and 0.36 of the windows in any strain, suggesting that gene flow between lineages is common in the wild (Figure 4C). Significant proportions of windows derived from the CN VIII, NA & Japan, Huangjiu & Baijiu and Beer1 & Bread clades were also observed (Figure 4C). Wild isolates with ≥0.1 genomic contribution from non-commercial wine lineages were correlated with phylogenetic (Figure 4A, Figure 4B) and PCA (Sup Figure 8) outliers to resolve four pseudo-populations; one population containing 58 samples with mixed Wine-Beer2 ancestry (AdB1) and three candidate admixed populations (AdEA1, AdEA2 and AdEA3), which contained windows with the closest *d*_*a*_ to modern East Asian lineages (Figure 4C). Of these East Asian influenced isolates, the AdEA1 population was the most common containing 33 samples isolated at a single location (W5) in two different grape variety ferments (Shiraz, Chardonnay) across 3 years (2016-2018). Population AdEA2 contained individuals (n = 5) from two locations (W4, W5) sampled in 2016 and 2018, whereas AdEA3 contained individuals (n = 2) from a single location (W5) and year (2018).

Admixture population allele frequency was then tested for allelic bias from the Beer2 or East Asia (Baijiu & Huangjiu, CN VIII and NA & Japan isolates) population that is absent from commercial and all other wild isolates (Wild*) using the 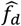 statistic^30^ (Figure 4C). Control tests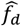 (Commercial, Wild*, East Asia; CN I) (95% CI: 0.0025 – 0.003; max: 0.016) and 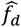 (Commercial, Wild*, East Asia, CN I) (95% CI: 0.0098 – 0.0011; max: 0.009) showed little deviation from the expected ∼0 value under a topology absent of admixture. However, tests using admixed populations displayed increased 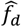 (Figure 4C): 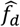 (Commercial, AdB1, Beer2; CN I) (95% CI: 0.0155 – 0.0182, max: 0.086), 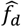 (Commercial, AdEA1, East Asia; CN I) (95% CI: 0.009 – 0.012; max: 0.176), 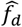 (Commercial, AdEA2, East Asia; CN I) (95% CI: 0.008 – 0.009; max: 0.1) and 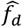 (Commercial, AdEA3, East Asia; CN I) (95% CI: 0.027 – 0.033, max: 0.21) suggesting loci have introgressed from East Asian linages dissimilar from those previously sequenced.

A drift tree was then constructed with the admixed populations separated from the bulk of the wild samples, reconstructing the demographic history of *S. cerevisiae* strains while accounting for between-clade gene flow (Figure 4D). Iterating through migration edge values 1-10 (Sup Figure 13) revealed widespread admixture between clades, likely contributing to the phylogenetic inconsistency reported in previous work^3,4,6^. After removing known admixture events^4,18,19^, the three novel admixture events were confirmed as being from unknown source lineages most related to modern East Asian populations (Figure 4D). The source lineage for the AdEA1 admixture event was estimated to have diverged from a common ancestor of East Asian domesticated ferments and contributed 0.12 drift to the AdEA1 population (Figure 4D). AdEA2 and AdEA3 sources were derived from likely non-domesticated lineages related to, but not a part of, the NA & Japan clade; contributing 0.14 and 0.44 drift respectively (Figure 4D).

### Multiple admixed haplotypes of divergent origin are segregating in wild populations

To confirm the presence of East Asian admixture and identify the breakpoints of admixed loci, two complementary approaches were applied to 10kb windows of computationally phased genotypes: 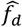 and local phylogenetic reconstruction. Signals of phylogenetic incongruence in these approaches were considered signals for admixture, confirming the presence of three distinct admixture events into wild isolates (Figure 5A, Sup Table 7). As the admixture source population is unsampled for each of the putative admixture events, local 10kb phylogenies were broken down into subtrees containing the test wild isolate, the closest sampled lineage to the putative admixture source, MO isolates and commercial wine isolates. Subtrees were considered incongruent if the wild isolate shared a MRCA with the closest sampled lineage (eg NA & Japan) while not in monophyly with MO and commercial isolates (Figure 5A i-iii).

**Figure 5:**
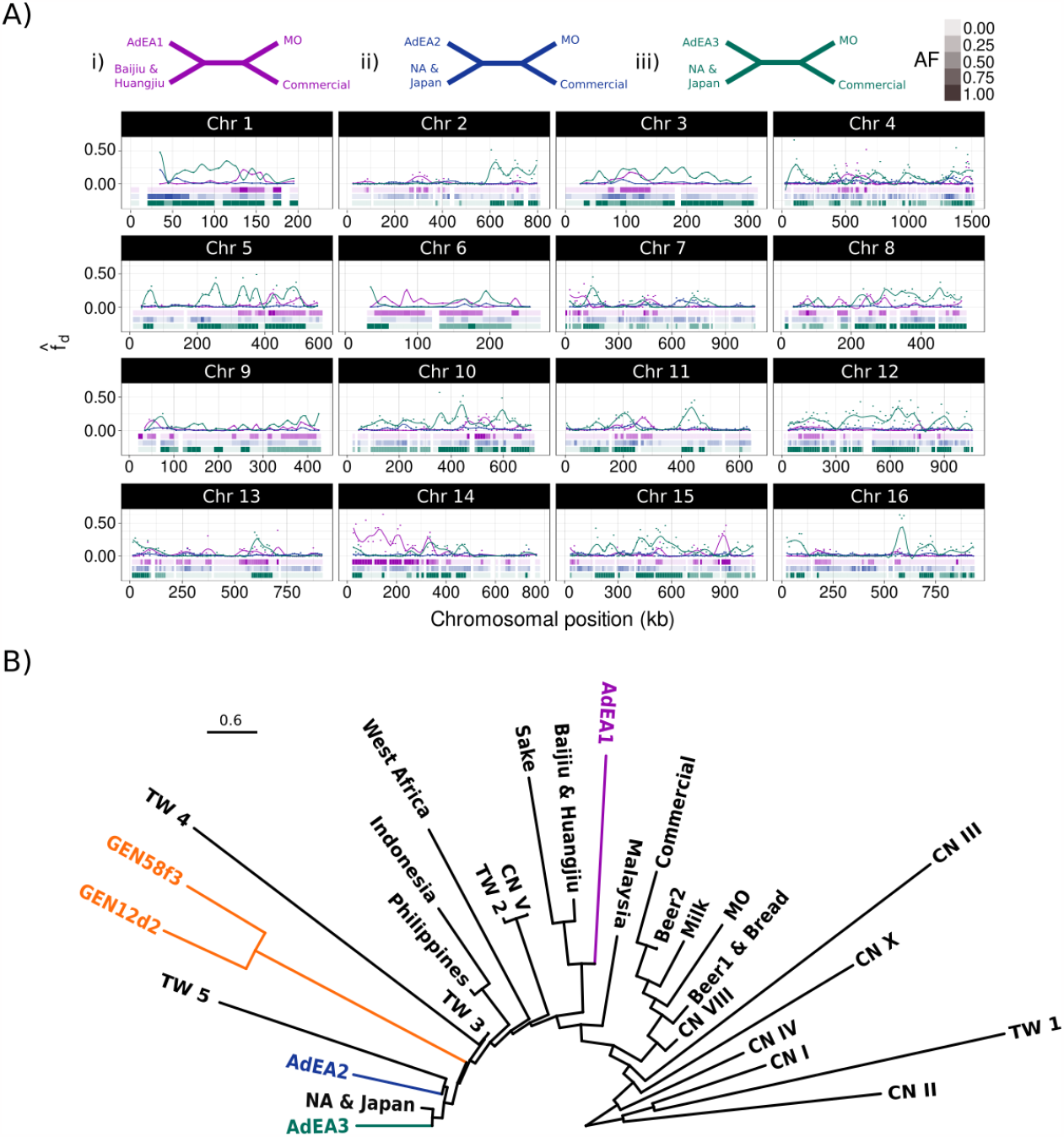
A) Genome wide windowed 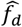 and local topology reconstruction identified fixed admixture loci in populations AdEA1 (purple), AdEA2 (blue) and AdEA3 (green). Each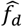 population was tested for their respective 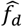 and subtree (i-iii) at each 10kb tiled window across the genome: Admixed East Asia 1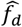 (Commercial, AdEA1; Baijiu&Huangjiu, CN I), ii) Admixed East Asia 2 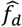 (Commercial, AdEA2; NA&Japan, CN I) and iii) Admixed East Asian 3 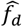 (Commercial, AdEA3; NA&Japan, CN I). Presence of admixture subtrees is shown as blocks below y=0 for each of the tested subtrees (i-iii) with transparency indicating allele frequency (AF) in the population. Completely colorless windows are due to the local trees being unresolved. B) Coalescent species tree reconstruction of fixed, consecutive admixture loci for AdEA1, AdEA2 and AdEA3 against a background of diverse lineages rooted using CN I, TW 1 and CN II populations. As no loci were fixed in AdEA2 a single homozygous diploid isolate from pop AdEA2 was used (Q-3_S92). *S. cerevisiae* isolates isolated from wild niches in Northern Australia (GEN12d2 and GEN58f3) are highlighted in orange. Scale is in coalescent units. An expanded tree with posterior probabilities can be found in Sup Figure 15.

Population AdEA1 showed clear signs of admixture from an unknown isolate basal to Baijiu & Huangjiu clade across 2.71 Mb (22.6% of genome) and fixed across 0.36 Mb (Figure 5A i). Due to the origin of the AdEA2 gene-flow event in the drift tree (Figure 4D) 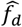 was not well resolved yet using the NA & Japan clade as the source population coupled with local phylogenetics clearly highlighted genetic breakpoints of the gene-flow event across 5.92 Mb (49% of genome), although none were fixed (Figure 5A ii). Population AdEA3 exhibited the largest total introgressed loci totalling 7.1 Mb (59% of genome) of which 3.7 Mb was fixed (Figure 5A iii).

Furthermore, the highly recombined and heterozygous nature of many of the introgressed loci from populations AdEA1 and AdEA2 (Figure 5A) suggests introgressed content is actively being removed from the standing variation. Yet some loci remained consistently fixed, for example, fixed loci in AdEA1 appeared stable over the 2016-2018 period where they were observed (Sup Figure 14), while other loci changed allele frequency between years (Sup Figure 14).

Phylogenetic reconstruction of homozygous admixture loci against a background of diverse populations, including recently identified Taiwanese wild populations^6^ along with further Chinese^3^ and South East Asian^8^ isolates confirmed they different from known *S. cerevisiae* lineages (Figure 5B, Sup Figure 15) and derive near estimated migration edges in the drift tree (Figure 4D). Population AdEA1 was derived from an unknown source lineage basal to East Asian domesticated yeast with high posterior support (Figure 5B, Sup Figure 15). Topology further revealed that AdEA2, AdEA3, and NA&Japan may be derived from a common ancestor, though support for these nodes was lower (Figure 5B, Sup Figure 15).

### Non-European derived populations exist in Australian wild niches

To determine if the donor lineages for admixture events AdEA1, AdEA2, and AdEA3 exist in wild ecological niches within Australia, an existing set of *S. cerevisiae* genome sequences isolated from environmental samples (data not shown) were screened for potential ancestry with the introgressed regions. Two related isolates were identified (GEN12d2 and GEN58f3), that were originally collected in Northern Australia, which displayed likely non-domesticated, non-european ancestry. GEN12d2 and GEN58f3 were highly homozygous displaying only 0.062% and 0.066% heterozygosity across the entire genome. Although genetic distance was low between the two isolates, some structural variation was identified, suggesting these are likely descended from the same individual. Phylogenetic placement of GEN12d2 and GEN58f3 across admixture loci revealed they share a most recent common ancestor with AdEA2, AdEA3 and NA & Japan isolates suggesting they may derive from the donor lineage of the AdEA2 and AdEA3 introgression events (Figure 5B). Furthermore, phylogenetic reconstruction using genome wide single copy orthologs against a background of phylogenetically diverse publicly available genomes (Sup Table 9) revealed that both GEN12d2 and GEN58f3, diverge from a common ancestor near the NA & Japan clade further supporting their non-domesticated, non-European origin (Sup Figure 16).

## Discussion

Spontaneous grape fermentations derive *S. cerevisiae* yeast from the surrounding environment, rather than starter cultures. Yet, little is known about the genetic diversity present within these natural populations. In this study we investigated the genomic landscape and architecture of wild yeast isolated from spontaneous grape fermentations in Australia. While it was found that Australian wild yeast were derived from highly recombined mixtures of multiple distinct commercial lineages, polysomic events and inter-clade admixture from Beer2 and unknown non-domesticated lineages potentially of Oceanic origin were found to be common.

Whole genome and chromosome duplication is a key strategy *S. cerevisiae* utilizes to rapidly adapt to novel stressors^31-35^. Due to *S. cerevisiae*’s selection for the fermentative environment, commercial wine strains may not be fit in the natural environment and need to adapt to survive. We found that whole genome duplications were observed at similar rates between wild yeast and commercial isolates, yet chromosomal duplications (aneuploidies) were far more common. Previous work^26^ found a negative correlation between chromosome length and aneuploid frequency. We recapitulate this result, however, relative frequency of chromosome polysomy was different between our study and previous work^3,8,36,37^ which may be due to unique stressors of the fermentative to wild to fermentative environmental transfer. We found that gain of chromosomes 1, 3, and 9 were common, agreeing with past studies^8,26,37^, where the increased copy number of these chromosomes has been implicated in responding to drug^33^, temperature ^38^, and environmental^26,39^ stress. Yet, chromosome 6 aneuploidy was also observed in many wild isolates despite being infrequent in past investigations^3,8,36,37^ and evidence suggesting chromosome 6 polysomy decreases viability^27,40,41^. However, chromosome 6 aneuploidy has been shown to be beneficial as a rescue mechanism for the compensation of deleterious mutations^31^. Furthermore, the observed frequency of chromosome 6 gain is likely possible due to its co-occurrence with chromosome 1 polysomy in all isolates, which has been previously suggested to increase its viability^41^. Taken together the results suggest that the high levels of chromosome 6 polysomy observed in spontaneous fermentations may confer an important fitness benefit absent from artificial environments.

Previous work^8^ suggested that wild *S. cerevisiae* strains show higher rates of heterozygosity than their commercial counterparts. In contrast, we found that commercial isolates had higher heterozygosity than wild samples, despite there being clear signs of sexual reproduction between wild isolates. One explanation for this conflicting observation comes from a genetic study of yeast derived from spontaneous wine fermentations, whereby heterozygous isolates transitioned into homozygous diploids through genome renewal^28^. This may have caused us to underestimate the rate of heterozygosity, but not nucleotide diversity, in wild isolates and future studies should investigate heterozygosity in environmental isolates.

Unlike beer, where fermentations are conducted year-round, with yeast reused across multiple ferments through pitching, wine production is seasonal and requires wine yeast to find suitable refuge between years. This forces *S. cerevisiae* to escape into the broader environmen^t4,22,23^, where they are subjected to novel selection pressures^15,42^. As wine strains exhibit decreased interbreeding though low spore viability^3,43,^ we expected wild yeast genomes would be made up of haplotypes derived from recombined commercial isolates. Yet, we observed evidence that wild populations contain both near-clonal commercial and highly mosaic isolates with few related commercial lineages contributing most loci. Though we are unable to determine if this occurred through biotic (e.g. selection for higher sporulation rates) or abiotic (e.g. increased usage) means, a clear conclusion is apparent: use of commercial strains in high frequency has a drastic effect on wild yeast diversity. This may be especially pertinent to countries, such as Europe and Asia, where native, divergent *S. cerevisiae* populations are widespread, potentially causing the wild standing variation to be overrun with few highly used commercial lineages through drift.

Transference of commercial isolates into the natural environment also provided the opportunity to investigate if gene flow has occurred between wild isolates and other *S. cerevisiae* genetic clades^4^. Admixture between *S. cerevisiae* clades^4^ has been an important source of adaptive novelty resulting in the origin of genetically and phenotypically distinct strains such as the Beer2 clade^18^. We observed three distinct admixture events between diverse non-domesticated isolates and wild Australian samples, which may be driving strain diversity. Phylogenetic reconstruction of each admixed locus identified that gene-flow was unique from those previously observed in Beer and Lager/Ale^18,19.^ Investigation into heterozygosity around admixed loci suggested that backcrossing with wild isolates is ongoing at most admixed loci. Yet, many loci have been retained across all years, despite others being removed, which may represent adaptive introgression, a phenomenon that has been observed in many species, providing novel genetic variation to respond to selection^44-46^.

Oceania is thought to lack native *S. cerevisiae* populations, only being described after wine and beer production began in the late 19^th^ century^47^. In contrast, evidence exists for Indigenous Australians carrying out fermentation utilizing non-*Saccharomyces* yeast lineages^48^. Presence of admixed samples presents a clear conundrum - where did the source lineages originate and how did they come to be present in Australia? The simplest explanation is that domesticated Asian yeast were introduced through fermentation of Sake and Huangjiu in Australia. Yet, local phylogenetic reconstruction revealed that branches leading to the three admixture events all occur before the most recent common ancestor of domesticated Sake, Baijiu, and Huangjiu suggesting donor lineages diverged pre-domestication. Endemic lineages of *S cerevisiae* are widespread across Southeast Asia^4,8^ and recent work isolated an individual from the CN X lineage in New Caledonia^4^, an island in the South Pacific approx. 1,100 km east of Australia.

Although these are genetically dissimilar to either admixture source lineage, they collectively suggest that the dispersal history of non-domesticated lineages may be far more complex and widespread than previously thought. Further supporting this, two samples isolated from wild niches in Northern Australia share a common ancestor with the donor of the AdEA2 and AdEA3 introgression events, which were isolated from Southern Australia. The geographic separation of these isolates suggests wild pre-domestication lineages are widespread in Australian contemporary populations which may predate colonisation.

Taken together our results show that wild *S. cerevisiae* cultured from spontaneous fermentations are highly aneuploid and admixed, containing gene flow from multiple independent lineages that has been maintained at fixation across multiple years. Despite this, closely related commercial lineages contribute the majority of genetics to the assessed wild isolates, providing evidence for the drastic effect commercial monoculture can have on the genetic diversity of natural microorganism populations. Finally, our results show that non-European derived non-domesticated lineages of *S. cerevisiae* are present in Australian ecological niches and are actively admixing with feral isolates of commercial origin.

## Methods

### Yeast strain selection and sequencing

Extracts from commercial spontaneous fermentations sourced from wineries within Australia were plated on WL media containing 25 µg/mL chloramphenicol and then arrayed on YPD media +25 µg/mL chloramphenicol in 96-well format using a PIXL automated colony-picker (Singer Instruments) or by hand. Individual colonies were identified using high-throughput ITS sequence analysis^48^. Isolates identified as *Saccharomyces cerevisiae* were cultured in liquid YPD, with DNA extracted using the Gentra Puregene Yeast/Bact. Kit (Qiagen). Whole-genome sequencing was performed using the Illumina Nextera XT library protocols and sequenced at 2 x 300bp read length on the MiSeq platform (Ramaciotti Centre for Functional Genomics, Randwick, Australia).

Short read data were input into jellyfish to construct 31-mer databases for each sample. Jellyfish dump was used to generate canonical 31-mers for each sample with a minimum number of observations greater than 4. Canonical 31-mers were queried against databases for each *Saccharomyces sensu stricto* species with an available reference genome (*S. cerevisiae*^24^, *S. paradoxus*^49^, *S. mikatae*^50^, *S. kudriavzevii^50^, S. jurei*^51^, *S. arboricous*^52^, *S. eubayanus*^53^, and *S. uvarum*^54^). This produced a presence/absence marker dataset for each sample against each *sensu stricto* reference genome. Markers for each sample were then filtered to remove markers that were present in more than one *sensu stricto* reference database, allowing for estimates of the proportion of genomic content from each *sensu stricto* species to be calculated.

Strains GEN12d2 and GEN58f3 were originally isolated from environmental samples (plant and feather sample respectively) collected from Bray Park, Queensland. Environmental samples were cultured in liquid YPD with 5% ethanol and 25 µg/mL chloramphenicol. Subsamples were then grown on WL + 25 µg/mL chloramphenicol and colonies were selected by hand and grown in YPD liquid. These selected individual colonies were identified using high-throughput ITS sequence analysis^48^. Isolates identified as *S. cerevisiae* were cultured in liquid YPD and DNA was extracted by lysis of protoplasts through digestion with zymolase and potassium acetate^55^. Nanopore sequencing libraries were prepared using the SQK-LSK112.24 kit and loaded into a FLO-MIN112 (R10.4) flow cell. Fast5 files were base called and demultiplexed using Guppy v6.4.8 (Oxford Nanopore Technologies, Oxford, UK) with the ‘sup’ model and a minimum quality score filtering of 7. A total of 91X and 119X coverage was obtained for strains GEN12d2 and GEN58f3, respectively.

### Quality control, mapping and genotyping

Multiple publicly available WGS resequencing datasets were used in this analysis containing commercial^16^ and non-wine^3,6-8^ isolates. SRA accessions can be found in Sup Table 1. Raw fastq data was quality controlled using fastqc (https://www.bioinformatics.babraham.ac.uk/projects/fastqc/) and ngsReports v2.2.3^56^. Samples that showed detectible levels of adapter contamination were trimmed using Trimmomatic v0.3.9^57^. Reads were then mapped to the S288C *S. cerevisiae* reference genome^24^ using NextGenMap v0.5.5^58^ under default settings to reduce reference mapping bias^59,60^. Aligned reads were then sorted, indexed and filtered (MAPQ>20) using SAMtools v1.6^61^ before being genotyped individually and merged by BCFtools v1.17^62^. Sample genotypes with GQ ≤ 20 or DP ≤ 6 were set to missing (./.) and then sites were removed if more than 80% of sample genotypes were missing. This produced the filtered genotype dataset used throughout the paper. To carry out local phylogenetics a second phased genotype set was made. This included additional samples from other diverse clades, see Sup Table 1 for accessions. Genotype phasing was carried out using SHAPEIT2 v2.r837^63^ under default settings without imputation. Both the filtered and phased VCF records were then converted to Genomic Data Structure (GDS) format using SeqArray v1.4.0^64^ for downstream analyses.

### Genome assembly and long-read variant calling

De-novo genome assemblies for *S. cerevisiae* strains GEN12d2 and GEN58f3 were performed using Canu v. 2.2^65^. A consensus sequence for each assembly was obtained using Medaka v1.7.3 (Oxford Nanopore Technologies, Oxford, UK). Reads were then mapped to the S288C genome using minimap2^66^ with the map-ont model. Variant calling was carried out against the S288C reference genome^24^ on reads with a minimum mapping quality of 20 using Medaka v1.7.3 (Oxford Nanopore Technologies, Oxford, UK) which was then filtered for GQ>30 using BCFtools v1.17^62^. Filtered long read genotypes were then incorporated into the phased genotype dataset using BCFtools v1.17^62^ merge under default settings.

Short read genome assembly was carried out for a subset of the MO samples and AdEA1, AdEA2, AdEA3 populations (Sup Table 8) using Velvet v1.2.10^67^ under default settings. The S288C gene annotations^68^ were then lifted over onto short read, longread and publicly available genome assemblies (Sup Table 8) using Liftoff v1.6.3^69^.

### Ploidy and aneuploidy estimation

Ploidy estimation was carried out using Smudgeplot v0.2.5^70^ for each commercial and wild sample with k=21. Per site read depth was extracted from the filtered genotype GDS using SeqArray v1.4.0^61^ and statistics calculated using the R programming language. To reduce bias from repetitive regions we calculated read depth only within 1386 single copy orthologs identified using BUSCO v4.1.2^71^ on the S228C annotation^68^. As individual sequencing reads are expected to sample each position in the genome randomly and in an independent manner^72^, aneuploidies were identified by comparing the distribution of per-site chromosomal read depth to the genome wide depth distribution (excluding the test chromosome) using a Mann-Whitney U-test. Chromosomes were considered to be aneuploidic if the rank biserial effect size was considered ‘large’ according to ^73^ ie rank biserial ≥ 0.5. Mann-Whitney tests were carried using the wilcoxon.test function in R and rank biserial effect size calculated using the effectsize v0.8.3 R package^74^. To test the behaviour of this method, a gain or loss aneuploidy event (Sup Figure 17) and segmental duplication of 0-50% (Sup Figure 18) of chromosome 6 were simulated and tested against a simulated genome with a background of 0-5 co-occurring aneuploidies (Sup Figure 17, Sup Figure 18; only 0 and 5 shown). Genome wide read depth distribution was sampled from a normal distribution with mean = 50 and σ = 10. The number of simulated sites for each chromosome were sampled according to the number of sites observed in the empirical dataset. Aneuploidies were simulated in the genome wide background by scaling read depth for 0-5 chromosomes to 3N (ie, 1.5x chromosome read depth), aneuploid chromosomes were selected from the empirically observed chromosome aneuploidy frequency reported by Gilchrist and Stelkens ^26^. Test chromosomes for aneuplodies were scaled by a factor iterating through -1 to 1 by 0.1 to simulate read depth bias (2N read depth + 2N read depth x scaling factor). Test chromosomes for duplications had a fraction of their length (−0.5 - 0.5) scaled by a factor of 1.5 (ie –0.5 has 1N for half the chromosome; 0.5 has 3N for half the chromosome). This showed that using a large rank biserial effect size (≥0.5) was robust at identifying true aneuploidies vs sequencing depth variation and segmental hemizygosity.

### Calculation of identity by state, coancestry coefficient and principal components

The filtered genotype GDS was passed to geaR v0.1.0 ^75^ to construct windows (10, 15, 20 and 25kb), with repeat regions excluded, and calculate nucleotide diversity across each wild genome to identify isolates with low levels (<0.005%) of heterozygosity. Identity by state for each pairwise comparison of commercial and wild wine yeast was calculated at the genome wide level with SNPrelate v1.32.0^76^. Genome wide calculation of the coancestry coefficient and Z scores were calculated by passing IBS values to the snpgdsCutTree function in SNPrelate v1.32.0^76^ with 50000 permutations and visualized using snpgdsDrawTree. The dendrogram was then manually inspected for clusters using a Z-score cut-off of 3 and an IBS of 0.99 to identify lineages. To identify commercial origin of wild loci, 10, 15, 20 and 25 kb tiled window analysis was then carried out by first partitioning non-repetitive variants into windows using gear v0.1.0^75^ then passed to SNPrelate v1.32.0^76^ for IBS calculation. Pairwise comparisons of wild vs commercial isolates were then carried out to identify the highest IBS values for each wild sample across windows at a 0.99 IBS cut-off. Genome wide contributions were then resolved into the lineages identified above to robustly determine commercial origin. Principal component analysis was carried out on the filtered genotype GDS file using SNPrelate v1.32.0^76^ and each principal component pair was visually inspected for stratification of wild isolates.

### Phylogenetics and population genomics

Linkage disequilibrium was calculated using the PopLDdecay tool v3.42^77^ under default settings. BUSCO v4.1.2^71^ was used to identify 1386 single copy orthologs in the S228C annotation^68^ and genotypes (including invariant sites) were extracted within SCOs using gear v0.1.0^75^ from the filtered genotype GDS. Gene trees were then reconstructed using IQ-TREE v2.2.2.3^78^ allowing for best model inference (-m MFP) (best-fit substitution models for each window can be found in Sup Table 9). Gene trees were then passed to ASTRAL-III v3.0^79^ to estimate the species phylogeny under default settings. ADMIXTURE v1.3.0^80^ was used to reconstruct the ancestral population structure at 2-5 values for the number of *a priori* populations (k). TreeMix v1.13^81^ was used to identify admixture from non-wine *S. cerevisiae* clade into outliers identified using PCA, topology and ADMIXTURE. Data was converted from GDS to tab separated allele frequency values as required by TreeMix using the R programming language. To further investigate these regions, within population/individual nucleotide diversity (Nei’s *π*)^29^, between population/individual absolute genetic distance (Nei’s *d*_*XY*_)^29^, and 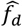 ^30^were calculated using the geaR v0.1.0 R package^75^ across 10kb tiled windows containing non-repetitive regions.

As the source population could not be utilised to identify incongruent topologies a subtree sorting method was implemented, inspired by the quartet subtree approach for comparing topologies^82^, to test local topology in the R programming language. Phased genotypes, that were not within repetitive regions, of all samples utilized in the whole genome tree (wild (n = 411), commercial (n = 169) and diverse strain origin (n = 91)) were extracted in 10kb windows and output to *fasta* with separate sequences for each haplotype using gear v0.1.0^75^. Maximum likelihood phylogenies were then reconstructed for each window using IQ-TREE v2.2.2.3^78^ utilizing ModelFinder Plus (-m MFP) to estimate the best fit substitution model for each alignment (best-fit substitution models for each window can be found in Sup Table 10). Local phylogenies were tested for incongruence by iterating across all admixed isolate haplotypes and extracting a subtree containing one admixed isolate haplotype, all individuals in the closest clade to the source of admixture (AdEA1:Baijiu&Huangjiu; AdEA2:NA&Japan; AdEA3:NA&Japan), Mediterranean oak isolates and commercial wine isolates with PhyTools v1.5^83^. As Mediterranean oak populations have been shown to share a common ancestor with European wine yeast after the dispersion of yeast out of Asia^7^, haplotypes that resolved in paraphyly with this MRCA can be considered incongruent. Local phylogenies with subtrees that were incongruent (i.e. at least one wild haplotype was both monophyletic with a source population and paraphyletic with all Commercial and Mediterranean Oak isolates) were treated as evidence for admixture.

Phylogenetic reconstruction of admixture source lineages was carried out by selecting loci with greater than or equal to three consecutive, fixed 10kb windows flagged by 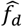 ^30^ and/or local topology as introgressed. As no loci were fixed in AdEA2 a single homozygous diploid isolate from pop AdEA2 was used (Q-3_S92). Windows at admixture breakpoints, ie the first and last 10kb windows, were then discarded as these may contain recombinant genotypes from both ancestries. Phylogenetic reconstruction was then carried out, incorporating the genotypes from the long read sequencing of GEN12d2 and GEN58f3, using IQ-TREE v2.2.23^78^ by selecting the best fit substitution model (-m MFP) for each window (best fit models can be found in Sup Table 11). All trees were then concatenated and passed to ASTRAL-III v3.0^79^ along with lineage assignments (Sup Table 1) to collapse clades into their respective lineages. As Astral-III can utilize local trees where not all leaves in the species tree are present, a single species tree was constructed incorporating all three East Asian admixture events even though they lacked homozygous overlapping introgression windows.

Whole genome phylogenetic reconstruction was carried out on genome assemblies by building individual gene trees from multiple sequence alignments carried out with MUSCLE v5.1^84^ for each of the 691 SCOs with acceptable gene alignments (≥5 isolates in alignment and ≥80% of the S288C gene length annotated) in IQ-TREE v2.2.2.3^78^ with model selection (-m MFP) (a list of models for each gene use can be found in Sup Table 12). Gene trees were then passed to Astral-III v2.2.2.3^79^ for species tree reconstruction under default settings.

## Data Availability

All raw short read and long read data and long read genome assemblies can be found in NCBI under project accession PRJNA980901.

## Acknowledgements

Special thanks to the students from Genesis Christian College for collecting the environmental samples from which GEN12d2 and GEN58f3 were isolated, as part of the Yeast Catchers Citizen Science Project. The AWRI, a member of the Wine Innovation Cluster in Adelaide, is supported by Australia’s grapegrowers and winemakers through their investment body Wine Australia with matching funds from the Australian Government. The Yeast Catchers project was supported by a Citizen Science grant from the Department of Industry, Science and Resources, Commonwealth of Australia. For genome sequencing the authors would like to thank the Ramaciotti Center for Genomics, which is funded through Bioplatforms Australia Pty Ltd (BPA), a National Collaborative Research Infrastructure Strategy (NCRIS).

## References

1. McGovern, P.E., Glusker, D.L., Exner, L.J. & Voigt, M.M. Neolithic resinated wine. Nature 381, 480–481 (1996).

2. McGovern, P.E. et al. Fermented beverages of pre-and proto-historic China. Proceedings of the National Academy of Sciences 101, 17593–17598 (2004).

3. Duan, S.-F. et al. The origin and adaptive evolution of domesticated populations of yeast from Far East Asia. Nature communications 9, 1–13 (2018).

4. Pontes, A., Hutzler, M., Brito, P.H. & Sampaio, J.P. Revisiting the Taxonomic Synonyms and Populations of Saccharomyces cerevisiae—Phylogeny, Phenotypes, Ecology and Domestication. Microorganisms 8, 903 (2020).

5. Cavalieri, D., McGovern, P.E., Hartl, D.L., Mortimer, R. & Polsinelli, M. Evidence for S. cerevisiae Fermentation in Ancient Wine. Journal of Molecular Evolution 57, S226–S232 (2003).

6. Lee, T.J. et al. Extensive sampling of Saccharomyces cerevisiae in Taiwan reveals ecology and evolution of predomesticated lineages. Genome Research 32, 864–877 (2022).

7. Almeida, P. et al. A population genomics insight into the Mediterranean origins of wine yeast domestication. Molecular Ecology 24, 5412–5427 (2015).

8. Peter, J. et al. Genome evolution across 1,011 Saccharomyces cerevisiae isolates. Nature 556, 339–344 (2018).

9. Di Maro, E., Ercolini, D. & Coppola, S. Yeast dynamics during spontaneous wine fermentation of the Catalanesca grape. International Journal of Food Microbiology 117, 201–210 (2007).

10. Granchi, L., Ganucci, D., Buscioni, G., Mangani, S. & Guerrini, S. The biodiversity of Saccharomyces cerevisiae in spontaneous wine fermentation: the occurrence and persistence of winery-strains. Fermentation 5, 86 (2019).

11. Sternes, P.R., Lee, D., Kutyna, D.R. & Borneman, A.R. A combined meta-barcoding and shotgun metagenomic analysis of spontaneous wine fermentation. Gigascience 6, gix040 (2017).

12. Callejon, R. et al. Volatile and sensory profile of organic red wines produced by different selected autochthonous and commercial Saccharomyces cerevisiae strains. Analytica Chimica Acta 660, 68–75 (2010).

13. Schvarczova, E., Stefanikova, J., Jankura, E. & Kolek, E. Selection of autochthonous Saccharomyces cerevisiae strains for production of typical Pinot Gris wines. Journal of Food &Nutrition Research 56(2017).

14. Alexandre, H. Wine yeast terroir: separating the wheat from the chaff—for an open debate. Microorganisms 8, 787 (2020).

15. Molinet, J. & Cubillos, F.A. Wild Yeast for the Future: Exploring the Use of Wild Strains for Wine and Beer Fermentation. Frontiers in Genetics 11(2020).

16. Borneman, A.R., Forgan, A.H., Kolouchova, R., Fraser, J.A. & Schmidt, S.A. Whole Genome Comparison Reveals High Levels of Inbreeding and Strain Redundancy Across the Spectrum of Commercial Wine Strains of Saccharomyces cerevisiae. G3 Genes|Genomes|Genetics 6, 957–971 (2016).

17. Marini, M.M. et al. The use of selected starter Saccharomyces cerevisiae strains to produce traditional and industrial cachaça: a comparative study. World Journal of Microbiology and Biotechnology 25, 235–242 (2009).

18. Fay, J.C. et al. A polyploid admixed origin of beer yeasts derived from European and Asian wine populations. PLOS Biology 17, e3000147 (2019).

19. Gallone, B. et al. Interspecific hybridization facilitates niche adaptation in beer yeast. Nature Ecology &Evolution 3, 1562–1575 (2019).

20. Hyma, K.E. & Fay, J.C. Mixing of vineyard and oak‐tree ecotypes of S accharomyces cerevisiae in N orth A merican vineyards. Molecular ecology 22, 2917–2930 (2013).

21. Tilakaratna, V. & Bensasson, D. Habitat predicts levels of genetic admixture in Saccharomyces cerevisiae. G3: Genes, Genomes, Genetics 7, 2919–2929 (2017).

22. Stefanini, I. et al. Social wasps are a Saccharomyces mating nest. Proceedings of the National Academy of Sciences 113, 2247–2251 (2016).

23. Zhang, H., Skelton, A., Gardner, R.C. & Goddard, M.R. Saccharomyces paradoxus and Saccharomyces cerevisiae reside on oak trees in New Zealand: evidence for migration from Europe and interspecies hybrids. FEMS Yeast Research 10, 941–947 (2010).

24. Mortimer, R.K. & Johnston, J.R. Genealogy of Principal Stains of the Yeast Genetic Stock Center. Genetics 113, 35–43 (1986).

25. Linder, R.A., Greco, J.P., Seidl, F., Matsui, T. & Ehrenreich, I.M. The Stress-Inducible Peroxidase TSA2 Underlies a Conditionally Beneficial Chromosomal Duplication in Saccharomyces cerevisiae. G3 Genes|Genomes|Genetics 7, 3177–3184 (2017).

26. Gilchrist, C. & Stelkens, R. Aneuploidy in yeast: Segregation error or adaptation mechanism? Yeast 36, 525–539 (2019).

27. Zhu, J., Pavelka, N., Bradford, W.D., Rancati, G. & Li, R. Karyotypic Determinants of Chromosome Instability in Aneuploid Budding Yeast. PLOS Genetics 8, e1002719 (2012).

28. Mortimer, R.K., Romano, P., Suzzi, G. & Polsinelli, M. Genome renewal: A new phenomenon revealed from a genetic study of 43 strains of Saccharomyces cerevisiae derived from natural fermentation of grape musts. Yeast 10, 1543–1552 (1994).

29. Nei, M. Molecular evolutionary genetics, (Columbia university press, 1987).

30. Martin, S.H., Davey, J.W. & Jiggins, C.D. Evaluating the Use of ABBA–BABA Statistics to Locate Introgressed Loci. Molecular Biology and Evolution 32, 244–257 (2014).

31. Filteau, M. et al. Evolutionary rescue by compensatory mutations is constrained by genomic and environmental backgrounds. Molecular Systems Biology 11, 832 (2015).

32. Chen, G., Bradford, W.D., Seidel, C.W. & Li, R. Hsp90 stress potentiates rapid cellular adaptation through induction of aneuploidy. Nature 482, 246–250 (2012).

33. Chen, G. et al. Targeting the adaptability of heterogeneous aneuploids. Cell 160, 771–784 (2015).

34. Selmecki, A.M., Dulmage, K., Cowen, L.E., Anderson, J.B. & Berman, J. Acquisition of aneuploidy provides increased fitness during the evolution of antifungal drug resistance. PLoS genetics 5, e1000705 (2009).

35. Selmecki, A.M. et al. Polyploidy can drive rapid adaptation in yeast. Nature 519, 349–352 (2015).

36. Gallone, B. et al. Domestication and divergence of Saccharomyces cerevisiae beer yeasts. Cell 166, 1397-1410. e16 (2016).

37. Sharp, N.P., Sandell, L., James, C.G. & Otto, S.P. The genome-wide rate and spectrum of spontaneous mutations differ between haploid and diploid yeast. Proceedings of the National Academy of Sciences 115, E5046–E5055 (2018).

38. Yona, A.H. et al. Chromosomal duplication is a transient evolutionary solution to stress. Proceedings of the National Academy of Sciences 109, 21010–21015 (2012).

39. Voordeckers, K. et al. Adaptation to high ethanol reveals complex evolutionary pathways. PLoS genetics 11, e1005635 (2015).

40. Liu, H., Krizek, J. & Bretscher, A. Construction of a GAL1-regulated yeast cDNA expression library and its application to the identification of genes whose overexpression causes lethality in yeast. Genetics 132, 665–673 (1992).

41. Torres, E.M. et al. Effects of aneuploidy on cellular physiology and cell division in haploid yeast. science 317, 916–924 (2007).

42. Kang, K. et al. Linking genetic, metabolic, and phenotypic diversity among Saccharomyces cerevisiae strains using multi-omics associations. Gigascience 8, giz015 (2019).

43. Puig, S., Querol, A., Barrio, E. &Pérez-Ortín, J. Mitotic Recombination and Genetic Changes in<i>Saccharomyces cerevisiae</i> during Wine Fermentation. Applied and Environmental Microbiology 66, 2057–2061 (2000).

44. Jones, M.R. et al. Adaptive introgression underlies polymorphic seasonal camouflage in snowshoe hares. Science 360, 1355–1358 (2018).

45. Pardo-Diaz, C. et al. Adaptive Introgression across Species Boundaries in Heliconius Butterflies. PLOS Genetics 8, e1002752 (2012).

46. Almeida, P., Barbosa, R., Bensasson, D., Gonçalves, P. & Sampaio, J.P. Adaptive divergence in wine yeasts and their wild relatives suggests a prominent role for introgressions and rapid evolution at noncoding sites. Molecular Ecology 26, 2167–2182 (2017).

47. Gayevskiy, V., Lee, S. & Goddard, M.R. European derived Saccharomyces cerevisiae colonisation of New Zealand vineyards aided by humans. FEMS Yeast Research 16(2016).

48. Varela, C. et al. A special drop: Characterising yeast isolates associated with fermented beverages produced by Australia’s indigenous peoples. Food Microbiology 112, 104216 (2023).

49. Liti, G., Barton, D.B. & Louis, E.J. Sequence diversity, reproductive isolation and species concepts in Saccharomyces. Genetics 174, 839–50 (2006).

50. Cliften, P. et al. Finding functional features in Saccharomyces genomes by phylogenetic footprinting. science 301, 71–76 (2003).

51. Naseeb, S. et al. Whole Genome Sequencing, de Novo Assembly and Phenotypic Profiling for the New Budding Yeast Species Saccharomyces jurei. G3 (Bethesda) 8, 2967–2977 (2018).

52. Liti, G. et al. High quality de novo sequencing and assembly of the Saccharomyces arboricolus genome. BMC Genomics 14, 69 (2013).

53. Baker, E. et al. The Genome Sequence of Saccharomyces eubayanus and the Domestication of Lager-Brewing Yeasts. Molecular Biology and Evolution 32, 2818–2831 (2015).

54. Vaughan Martini, A. & Kurtzman, C.P. Deoxyribonucleic acid relatedness among species of the genus Saccharomyces sensu stricto. International Journal of Systematic and Evolutionary Microbiology 35, 508–511 (1985).

55. Davis, R.W. et al. apid DNA isolations for enzymatic and hybridization analysis. In Methods in enzymology, Vol. 65 404–411 (Elsevier, 1980).

56. Ward, C.M., To, T.-H. & Pederson, S.M. ngsReports: a Bioconductor package for managing FastQC reports and other NGS related log files. Bioinformatics 36, 2587–2588 (2020).

57. Bolger, A.M., Lohse, M. & Usadel, B. Trimmomatic: a flexible trimmer for Illumina sequence data. Bioinformatics 30, 2114–2120 (2014).

58. Sedlazeck, F.J., Rescheneder, P. & von Haeseler, A. NextGenMap: fast and accurate read mapping in highly polymorphic genomes. Bioinformatics 29, 2790–2791 (2013).

59. Günther, T. & Nettelblad, C. The presence and impact of reference bias on population genomic studies of prehistoric human populations. PLOS Genetics 15, e1008302 (2019).

60. Ward, C.M. & Baxter, S.W. Assessing Genomic Admixture between Cryptic Plutella Moth Species following Secondary Contact. Genome Biology and Evolution 10, 2973–2985 (2018).

61. Li, H. et al. The Sequence Alignment/Map format and SAMtools. Bioinformatics 25, 2078–2079 (2009).

62. Li, H. A statistical framework for SNP calling, mutation discovery, association mapping and population genetical parameter estimation from sequencing data. Bioinformatics 27, 2987–2993 (2011).

63. Delaneau, O., Coulonges, C. & Zagury, J.-F. Shape-IT: new rapid and accurate algorithm for haplotype inference. BMC Bioinformatics 9, 540 (2008).

64. Zheng, X. et al. SeqArray—a storage-efficient high-performance data format for WGS variant calls. Bioinformatics 33, 2251–2257 (2017).

65. Koren, S. et al. Canu: scalable and accurate long-read assembly via adaptive k-mer weighting and repeat separation. Genome research 27, 722–736 (2017).

66. Li, H. Minimap2: pairwise alignment for nucleotide sequences. Bioinformatics 34, 3094–3100 (2018).

67. Zerbino, D.R. & Birney, E. Velvet: algorithms for de novo short read assembly using de Bruijn graphs. Genome research 18, 821–829 (2008).

68. Fisk, D.G. et al. Saccharomyces cerevisiae S288C genome annotation: a working hypothesis. Yeast 23, 857–65 (2006).

69. Shumate, A. & Salzberg, S.L. Liftoff: accurate mapping of gene annotations. Bioinformatics 37, 1639–1643 (2021).

70. Ranallo-Benavidez, T.R., Jaron, K.S. & Schatz, M.C. GenomeScope 2.0 and Smudgeplot for reference-free profiling of polyploid genomes. Nature Communications 11, 1432 (2020).

71. Simão, F.A., Waterhouse, R.M., Ioannidis, P., Kriventseva, E.V. & Zdobnov, E.M. BUSCO: assessing genome assembly and annotation completeness with single-copy orthologs. Bioinformatics 31, 3210–3212 (2015).

72. Lander, E.S. & Waterman, M.S. Genomic mapping by fingerprinting random clones: a mathematical analysis. Genomics 2, 231–9 (1988).

73. Cohen, J. Statistical power analysis for the behavioral sciences, (Academic press, 2013).

74. Ben-Shachar, M.S., Lüdecke, D. & Makowski, D. effectsize: Estimation of effect size indices and standardized parameters. Journal of Open Source Software 5, 2815 (2020).

75. Ward, C.M., Ludington, A.J., Breen, J. & Baxter, S.W. Genomic evolutionary analysis in R with geaR. bioRxiv, 2020.08.06.240754 (2020).

76. Zheng, X. et al. A high-performance computing toolset for relatedness and principal component analysis of SNP data. Bioinformatics 28, 3326–3328 (2012).

77. Zhang, C., Dong, S.S., Xu, J.Y., He, W.M. & Yang, T.L. PopLDdecay: a fast and effective tool for linkage disequilibrium decay analysis based on variant call format files. Bioinformatics 35, 1786–1788 (2019).

78. Nguyen, L.-T., Schmidt, H.A., von Haeseler, A. & Minh, B.Q. IQ-TREE: A Fast and Effective Stochastic Algorithm for Estimating Maximum-Likelihood Phylogenies. Molecular Biology and Evolution 32, 268–274 (2014).

79. Zhang, C., Rabiee, M., Sayyari, E. & Mirarab, S. ASTRAL-III: polynomial time species tree reconstruction from partially resolved gene trees. BMC Bioinformatics 19, 153 (2018).

80. Alexander, D.H., Novembre, J. & Lange, K. Fast model-based estimation of ancestry in unrelated individuals. Genome Res 19, 1655–64 (2009).

81. Pickrell, J.K. & Pritchard, J.K. Inference of Population Splits and Mixtures from Genome-Wide Allele Frequency Data. PLOS Genetics 8, e1002967 (2012).

82. Estabrook, G.F., McMorris, F. & Meacham, C.A. Comparison of undirected phylogenetic trees based on subtrees of four evolutionary units. Systematic Zoology 34, 193–200 (1985).

83. Revell, L.J. phytools: an R package for phylogenetic comparative biology (and other things). Methods in Ecology and Evolution 3, 217–223 (2012).

84. Edgar, R.C. MUSCLE: multiple sequence alignment with high accuracy and high throughput. Nucleic acids research 32, 1792–1797 (2004).

