## Supplementary Figures for "The genomic landscape of wild *Saccharomyces cerevisiae* is shaped by complex patterns of admixture, aneuploidy and recombination"

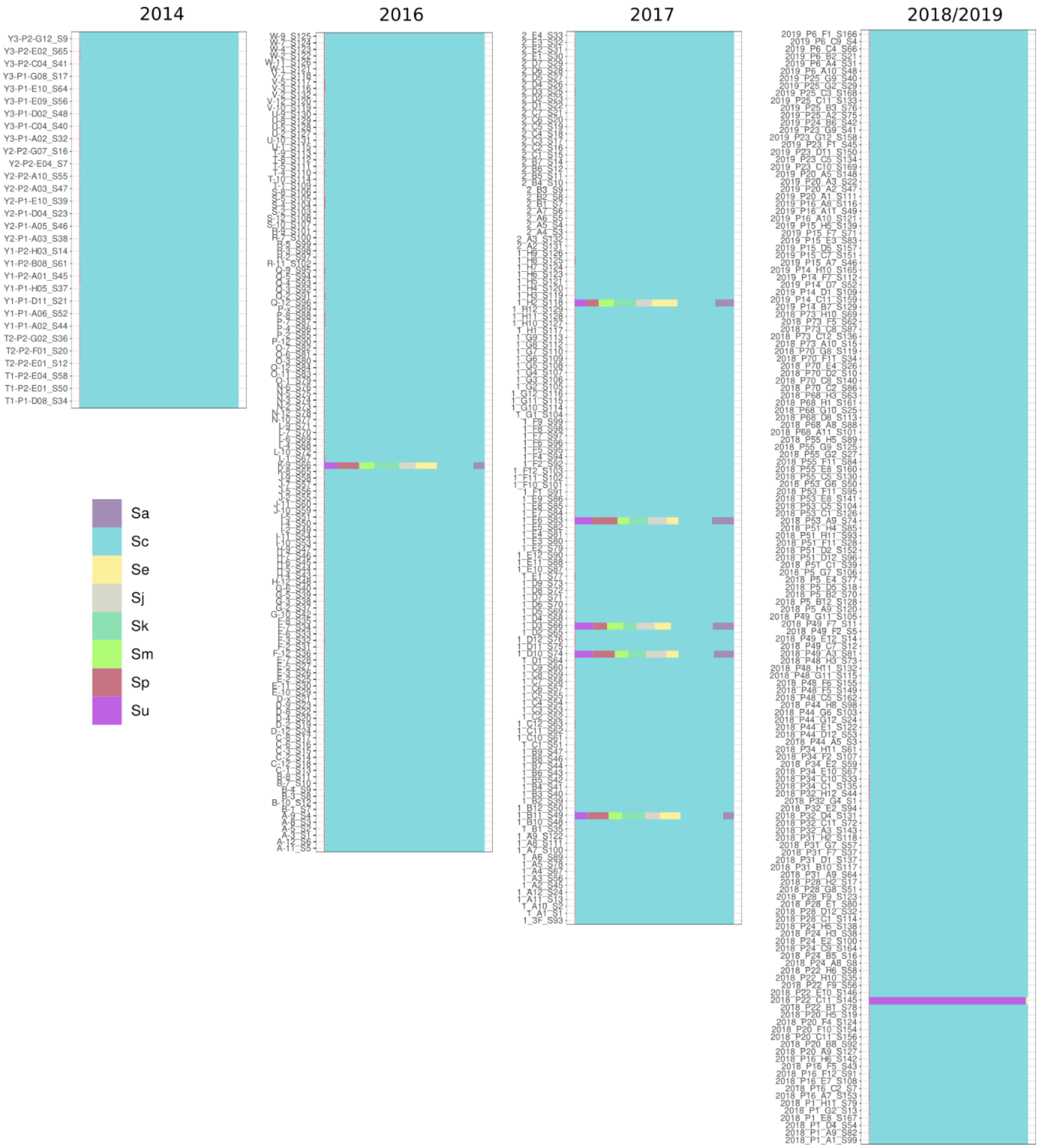


Sup Figure 1: Genome composition of wild isolates. Proportion of 21-mers with an exact match to one of the *sensu stricto Saccharomyces* species *S. arboricola* (Sa), *S. cerevisiae* (Sc), *S. eubayanus* (Se), *S. jurei* (Sj), *S. kudriavzevii* (Sk), *S. mikatae* (Sm), *S. paradoxus* (Sp), and *S. uvarum* (Su) for each wild isolate.

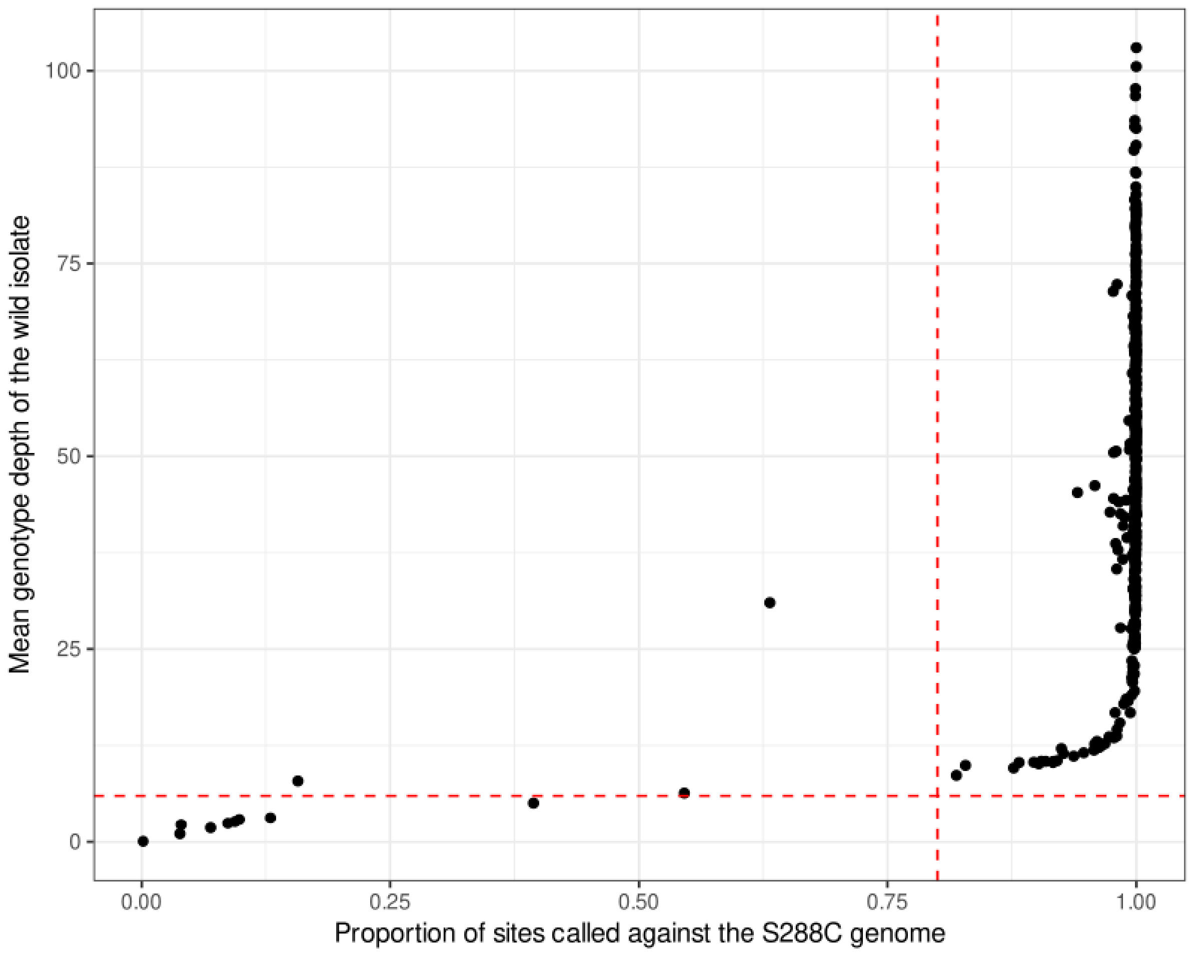


Sup Figure 2: Genome wide mean read depth mapped to the S288C genome against the proportion of the genome genotyped with red dashed lines showing the cutoffs for sample retention DP>5 and proportion > 0.8.

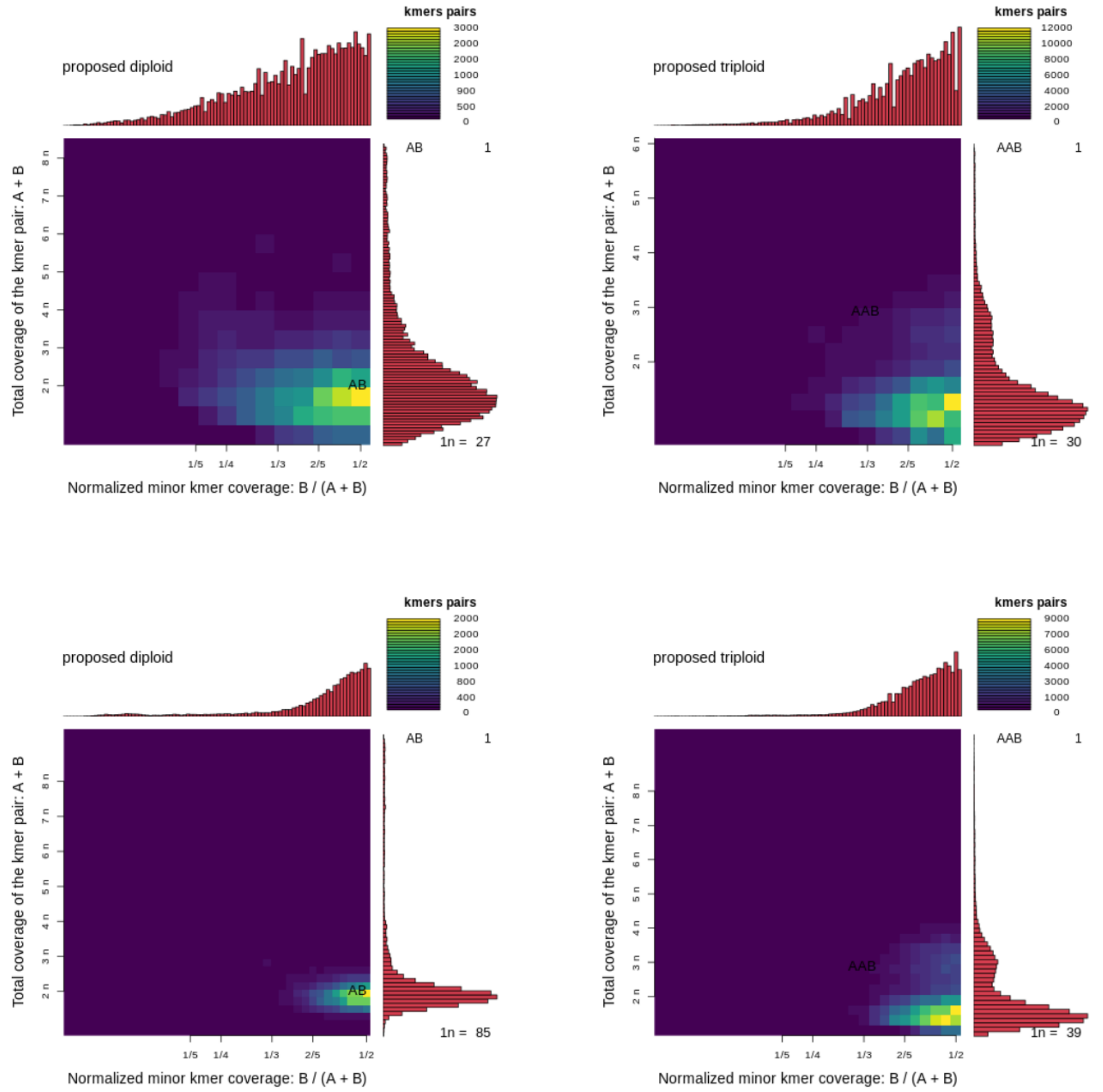


Sup Figure 3: Example of heatmaps generated by SmudgePlot for ploidy inference based on k-mers showing both a diploid and polyploid islate. Upper: Wild samples, Lower: Commercial samples

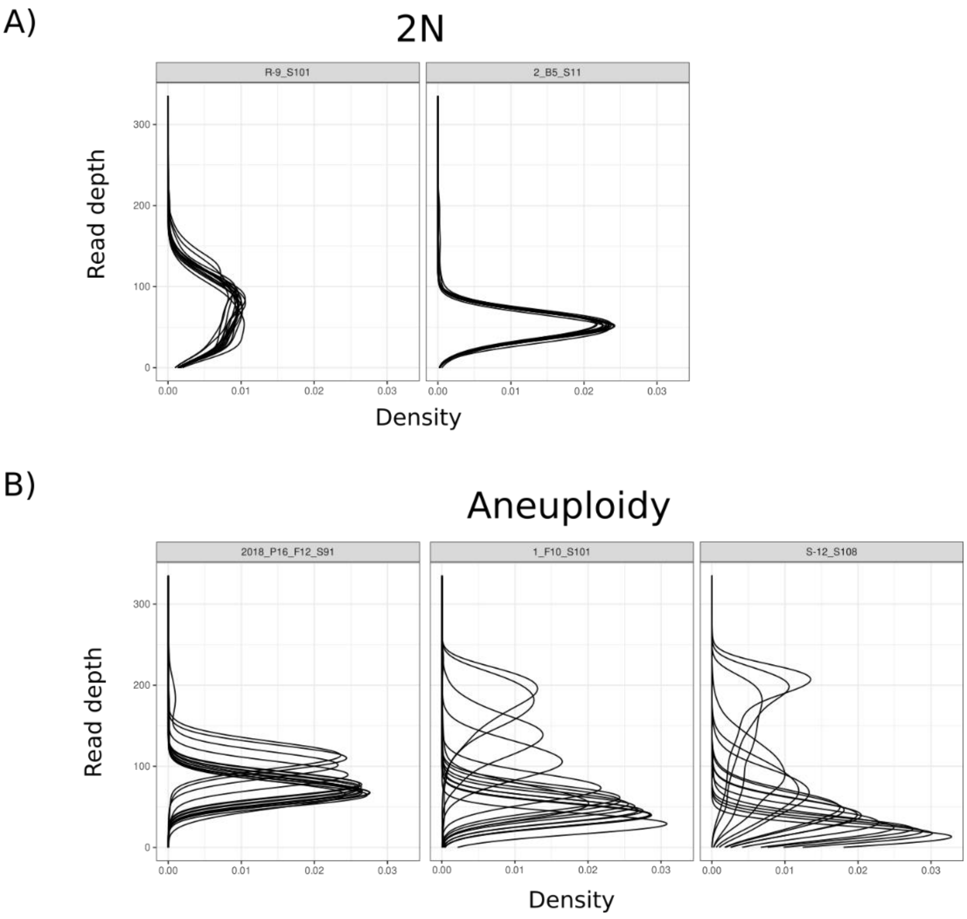


Sup Figure 4: Read depth density plots broken down by chromosome for the same samples shown in Figure 1. A) 2N. B) Aneuploids.

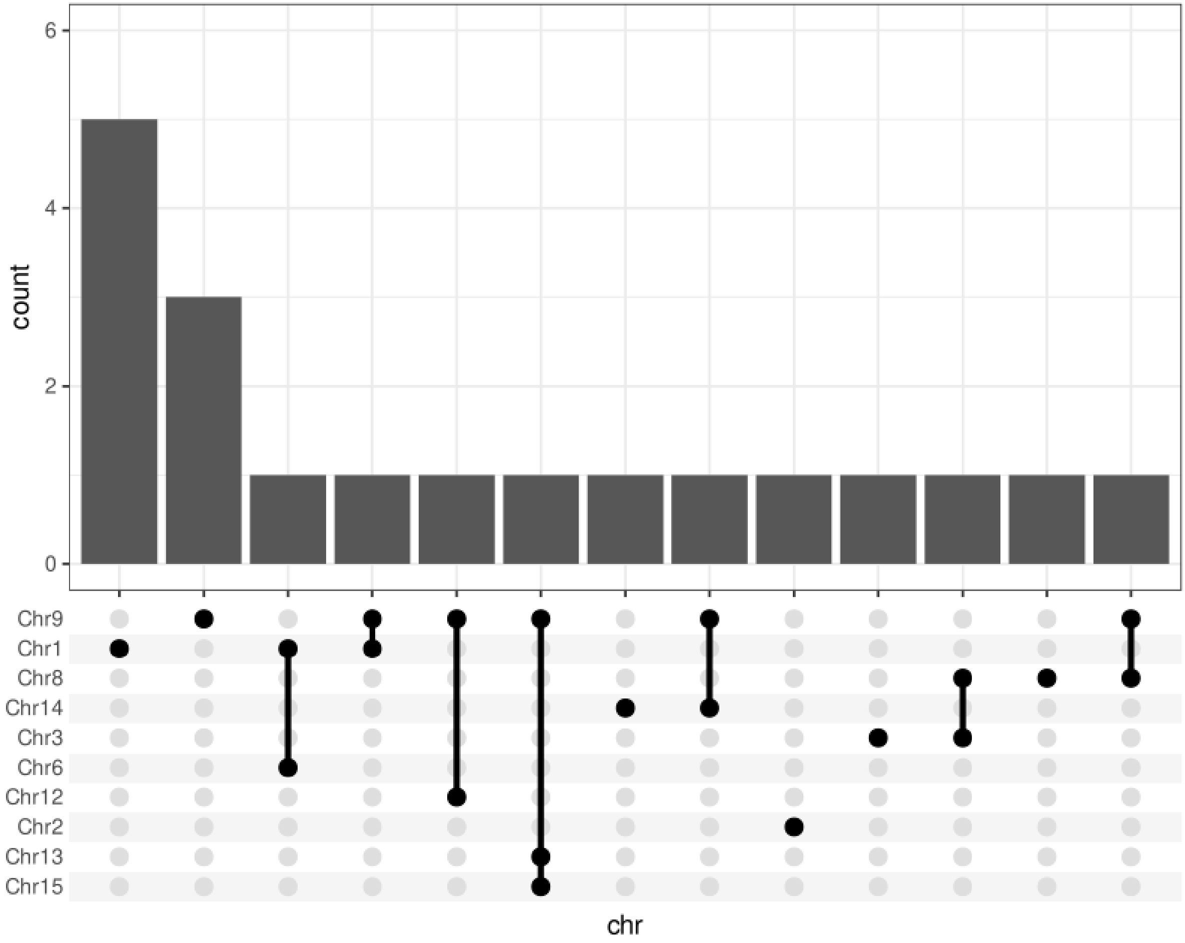


Sup Figure 5: Chromosome gain events observed in commercial isolates. Upset plot nodes represent aneuploidies that were present simultaneously in the same individual.

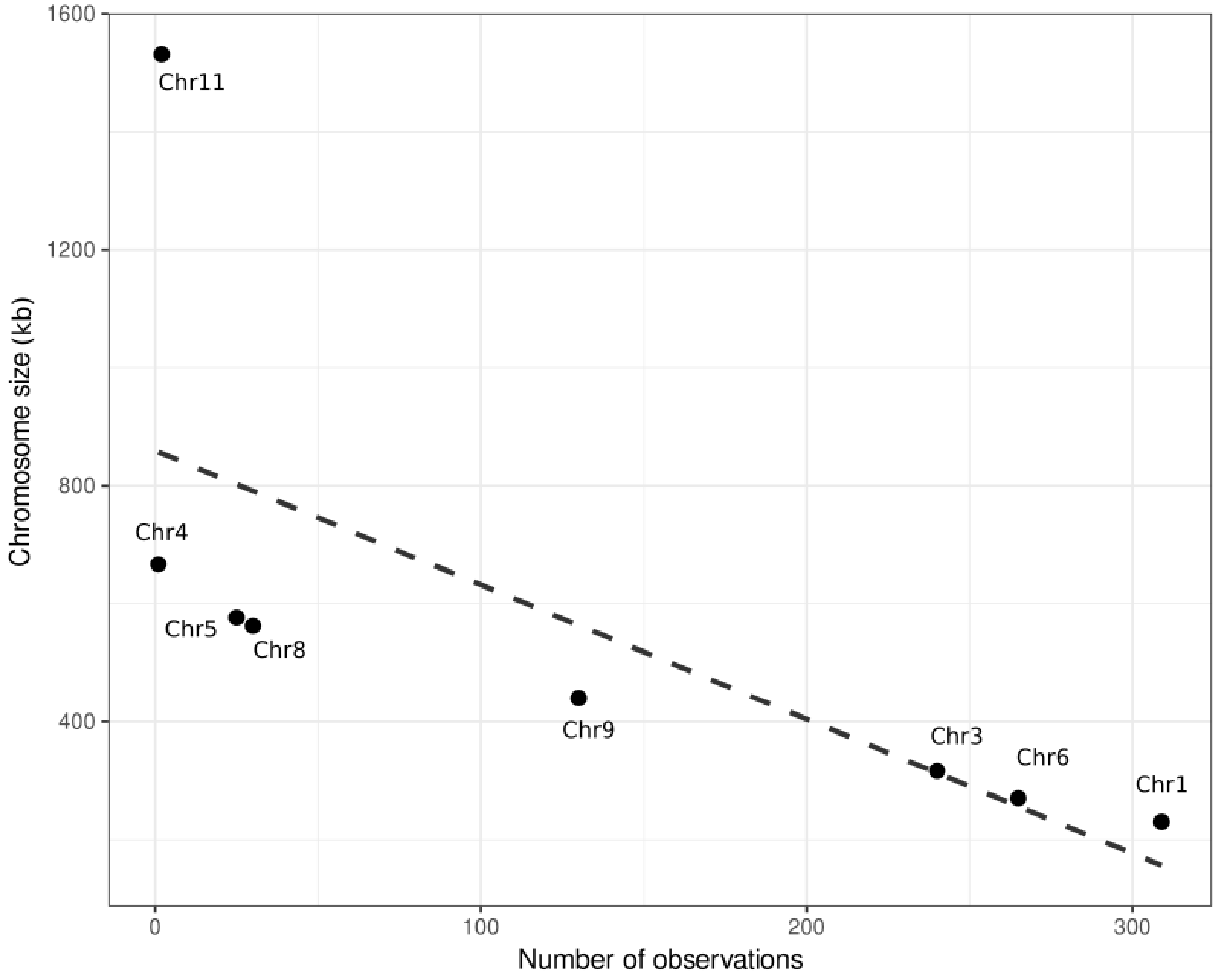


Sup Figure 6: Aneuploidies show a negative correlation with chromosome size. Linear model fit to the number of observations of aneuplodies for each chromosome across the entire Wild isolate pool against chromosome size.

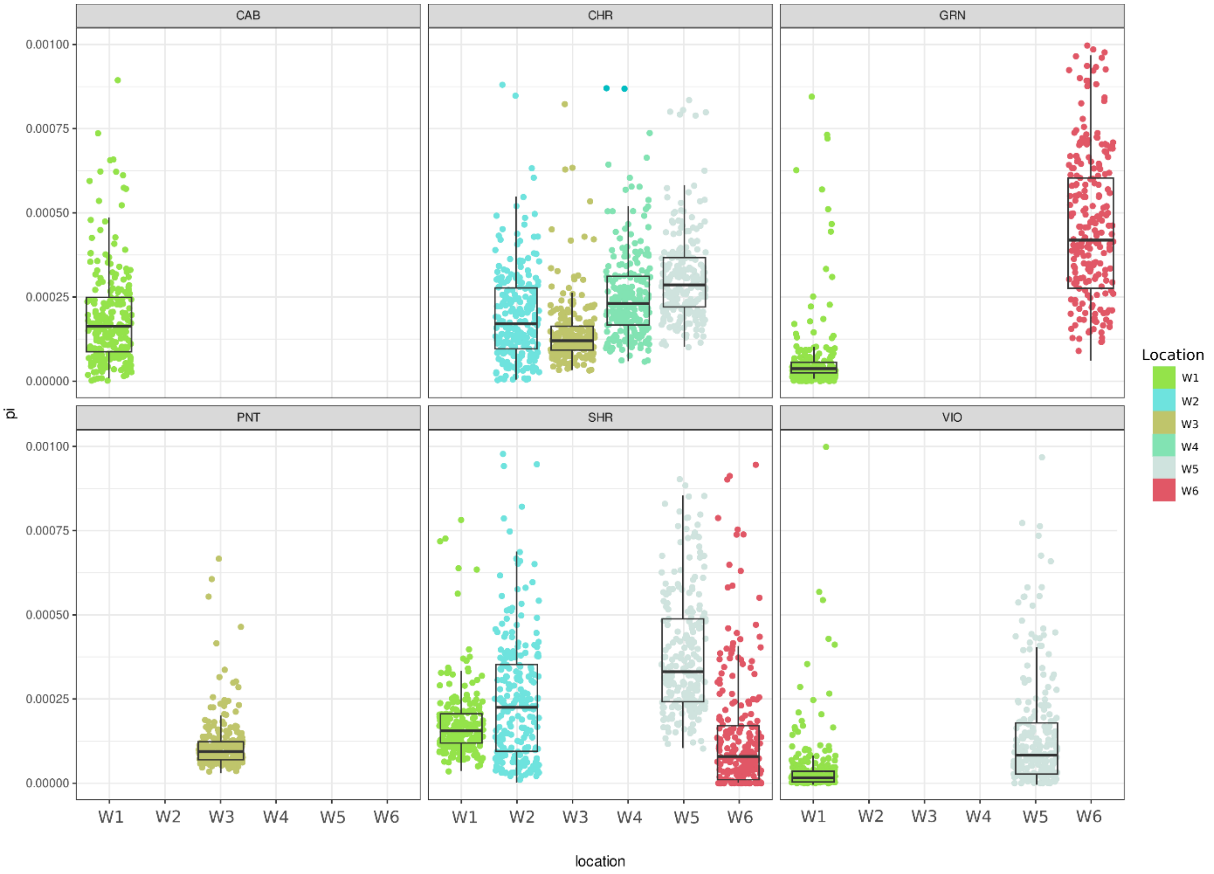


Sup Figure 7: Nucleotide diversity is different between locations when you control for grape variety. Jitter plot showing nucleotide diversity (pi) distributions calculated within 25kb windows across the genome, boxplots were overlaid with boxes showing the first and third inter quartile range (IQR) while whiskers extend to a maximum of 1.5 * IQR. VIO = Viognier, PNT = Pinot Noir, SHR = Shiraz, GRN = Grenache, CHR = Chardonnay, CAB = Cabernet Sauvignon.

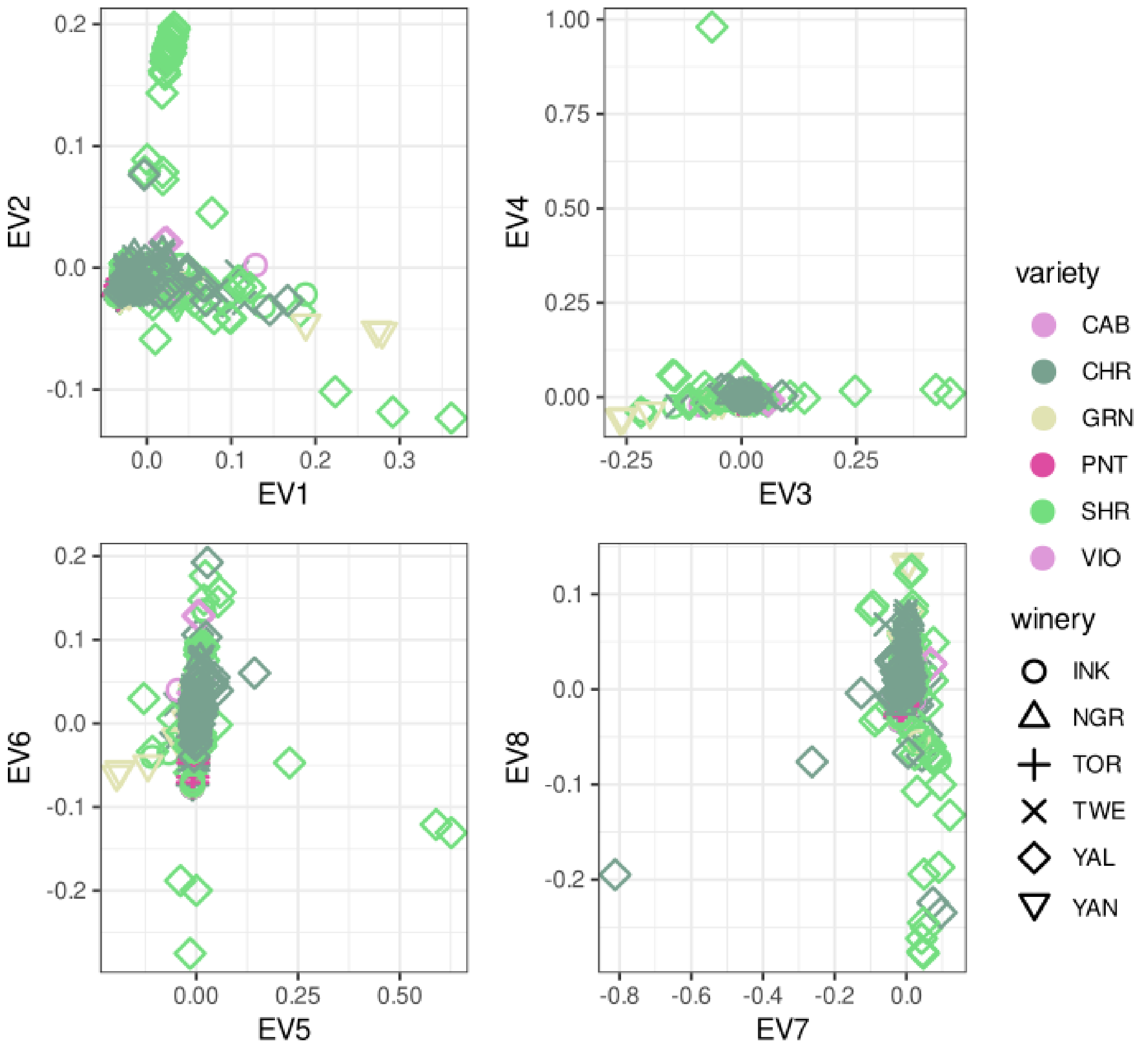


Sup Figure 8: Principal component analysis of wild isolates shows a lack of widespread population structure between locations and grape varieties across eigenvectors (EV) 1-8.

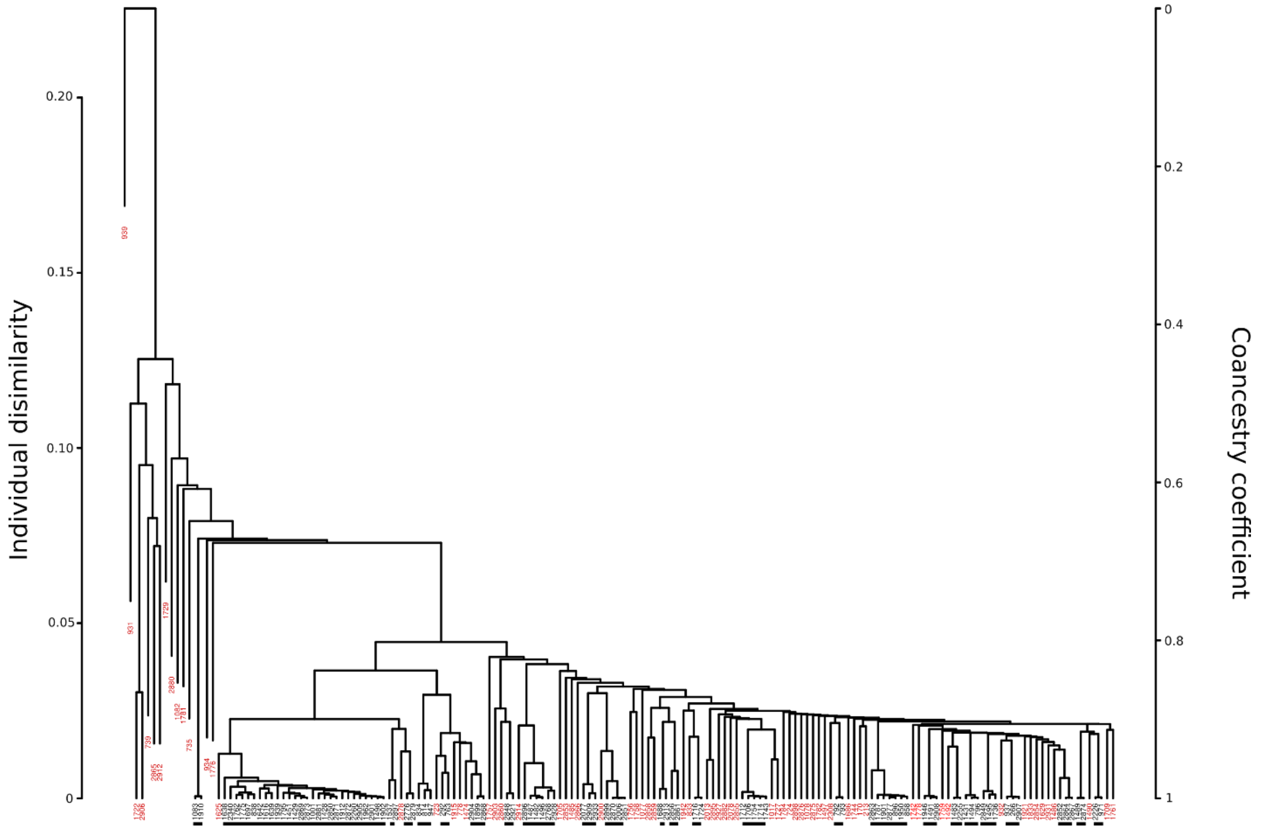


Sup Figure 9: Co-ancestry coefficient dendrogram for all commercial wine isolates with clack lines showing those isolates considered lineages for IBS analysis. Red isolates are unique isolates that do not form a lineage.

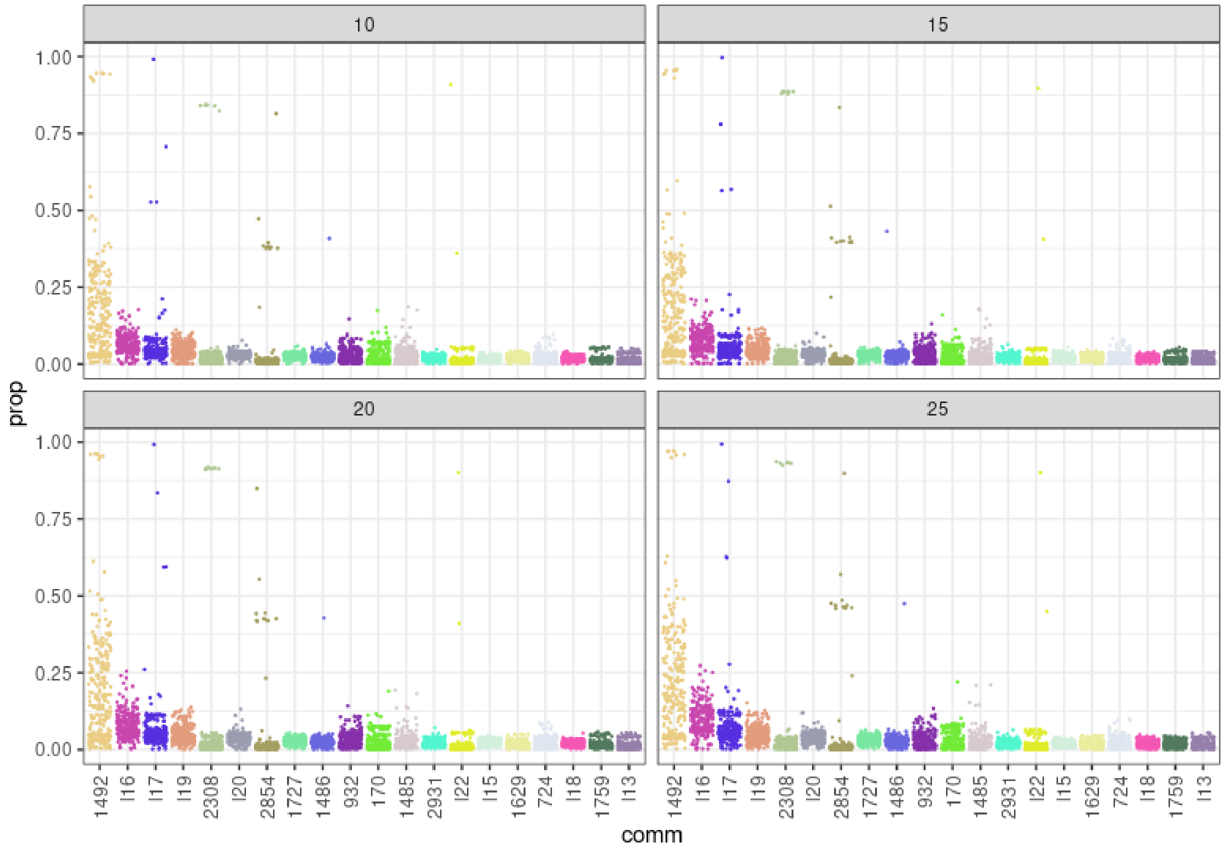


Sup Figure 10: The proportion of windows assigned to the top 20 lineages identified using co-ancestry analysis in Sup Figure 9. Four different widow sizes 10, 15, 20 and 25 kb to test for bias in window assignment based on window size.

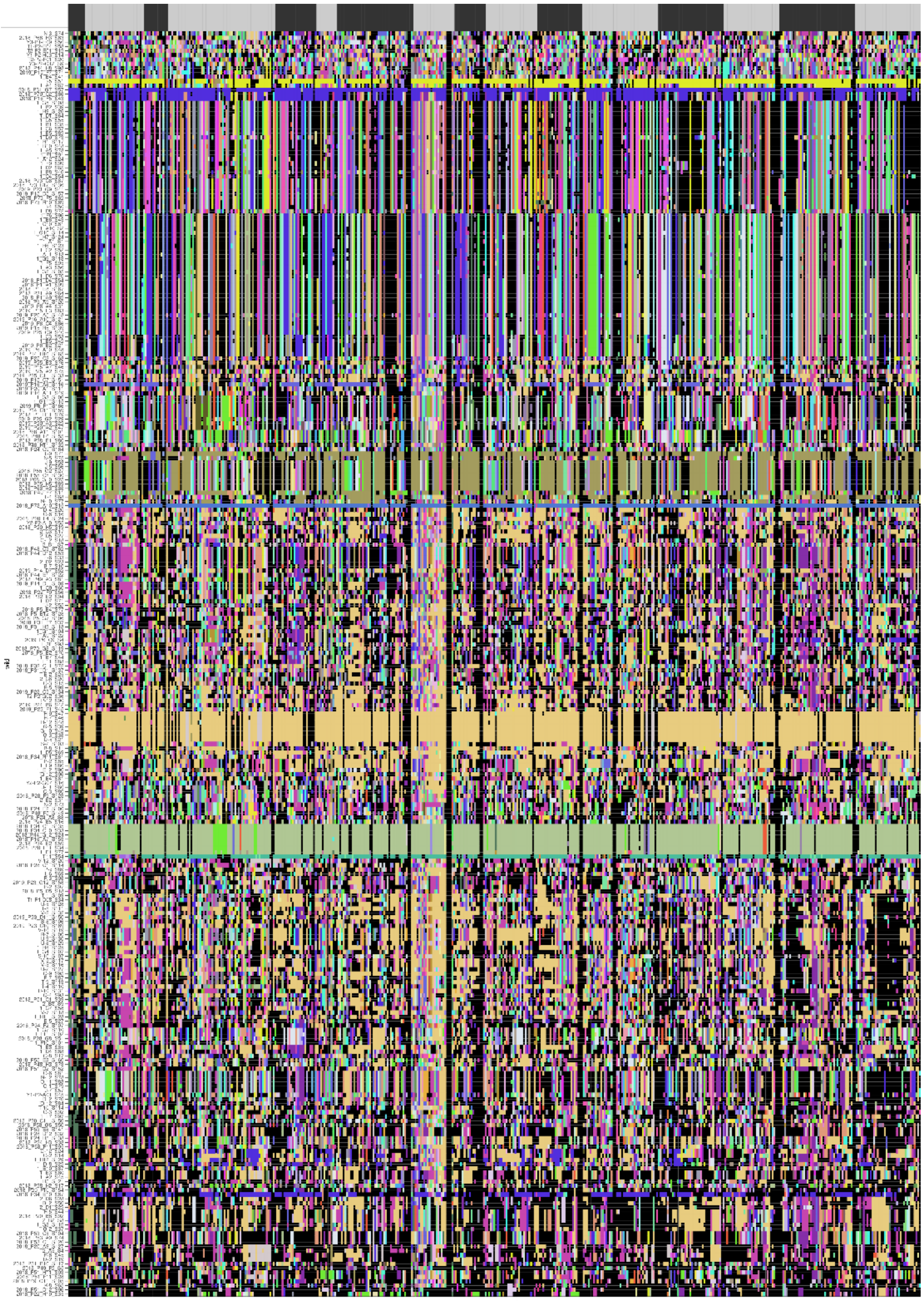

Sup Figure 11: Window-wise 25kb Identity by State calculated for all pairwise comparisons of commercial and homozygous diploid wild isolates. Colors indicate assigned commercial strain lineage of origin. Block-wise assignment to chromosomes can be found above the figure.

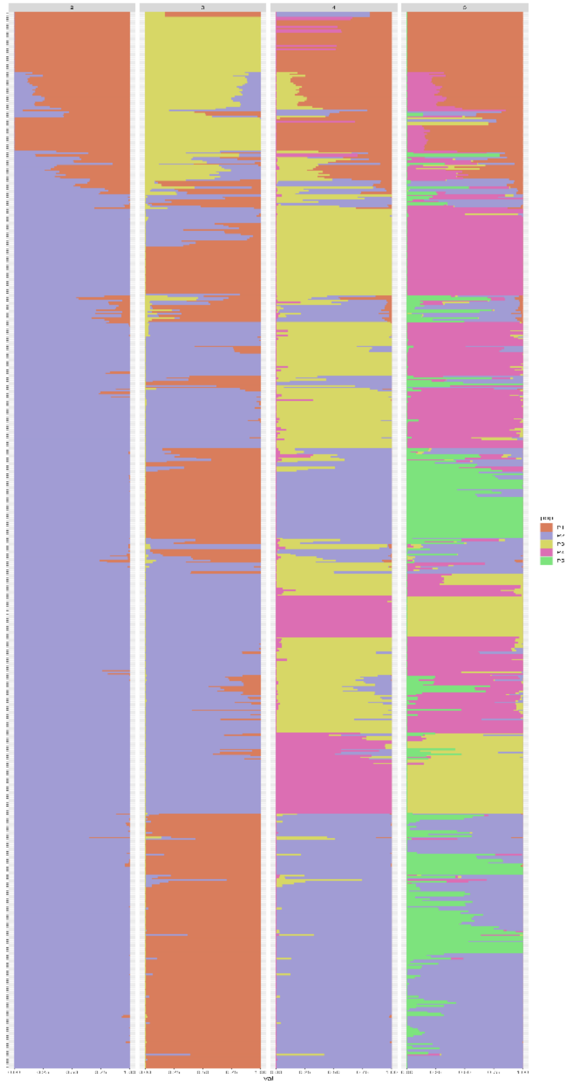

Sup Figure 12: ADMIXTURE ancestral population structure inference with *a priori* population number set from 2-5 ordered by their phylogenetic relationship shown in Figure 5A

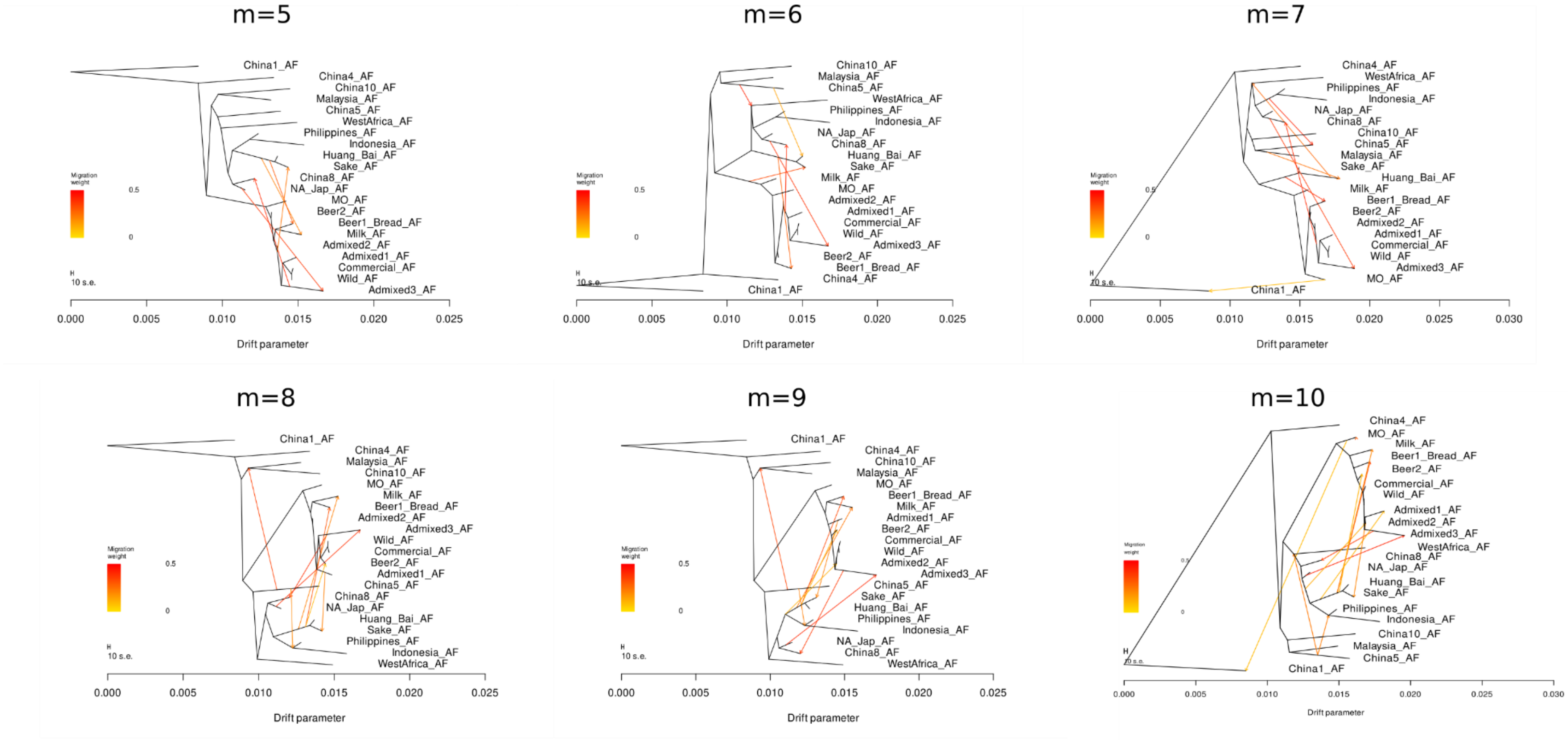


Sup Figure 13: TreeMix drift trees iterating through number of migration events (m) 1-10 reveals widespread admixture between genetic clades. Migration edges are coloured based on the estimated proportion of drift from source to target population.

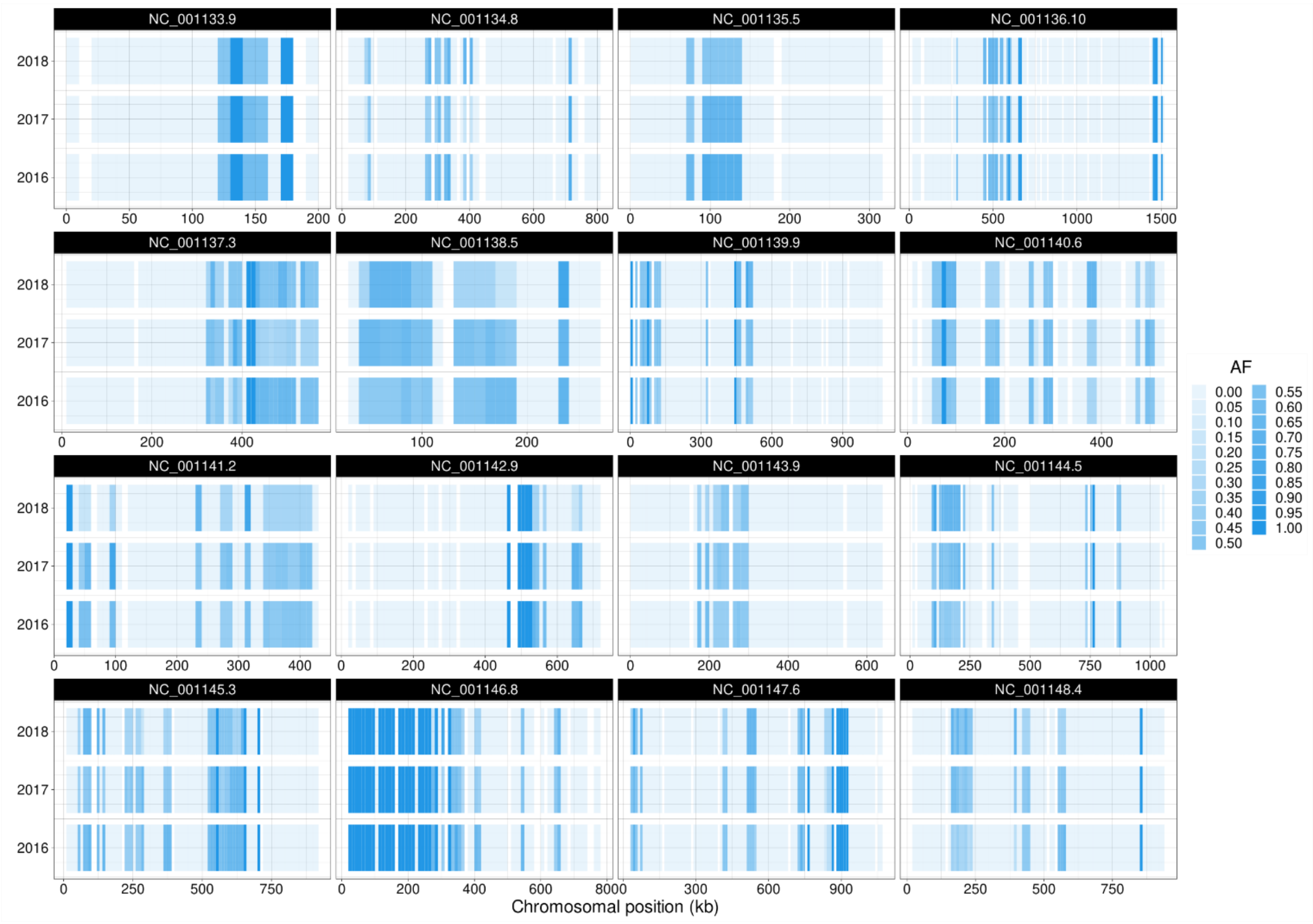


Sup Figure 14: 10kb windowed allele frequency of admixed loci in population AdEA1 across the three years it was observed 2016, 2017 and 2018. Blank windows are where the local topology was unresolved.

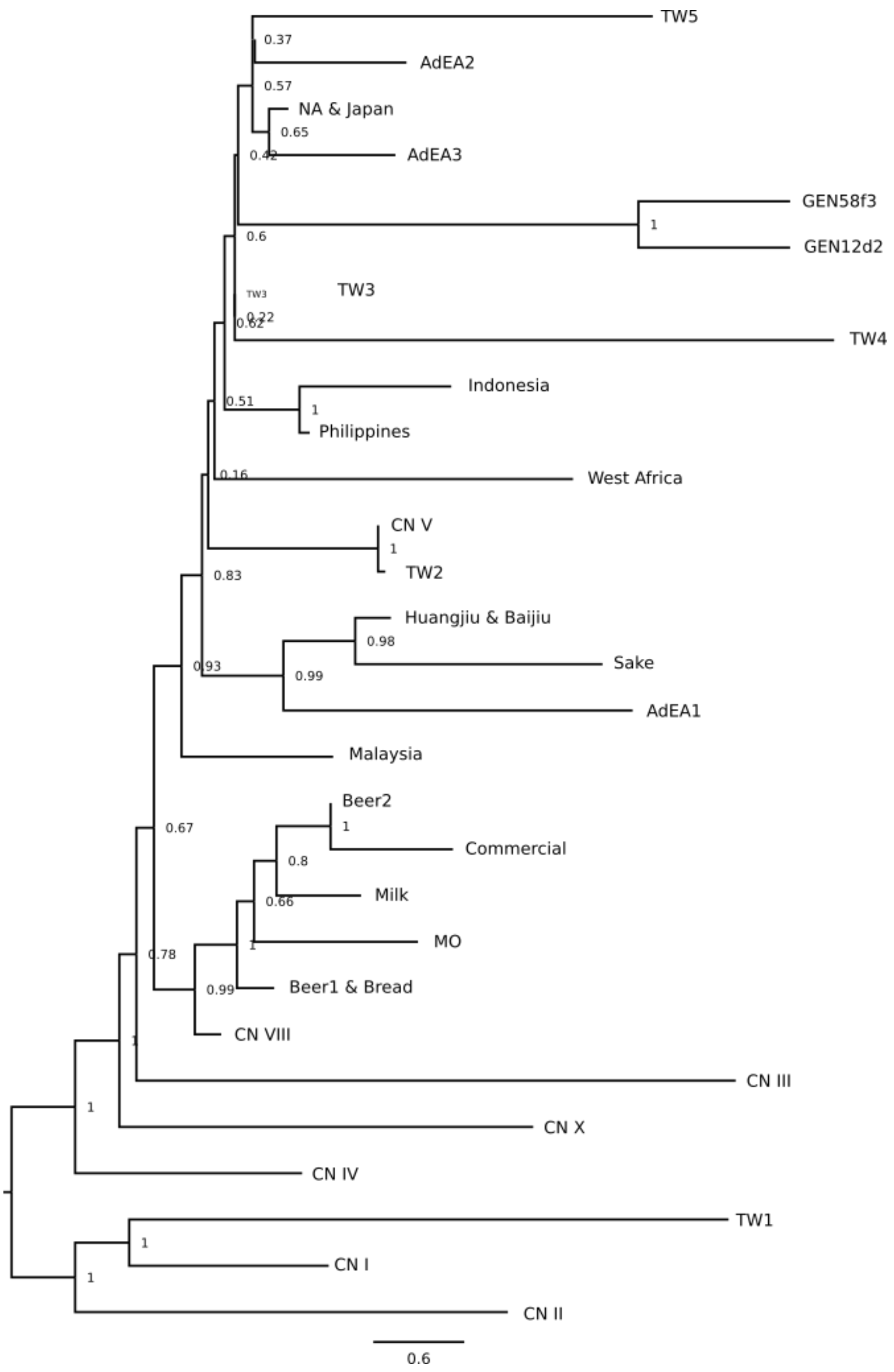


Sup Figure 15: Coalescent species tree reconstruction of diverse *S. cerevisiae* populations along with AdEA1, AdEA2 and AdEA3 using fixed admixture loci identified in Figure 5A. As no loci were fixed in AdEA2 a single homozygous diploid isolate from pop AdEA2 was used (Q-3_S92). Node posterior probability is shown. Scale is in coalescent units.

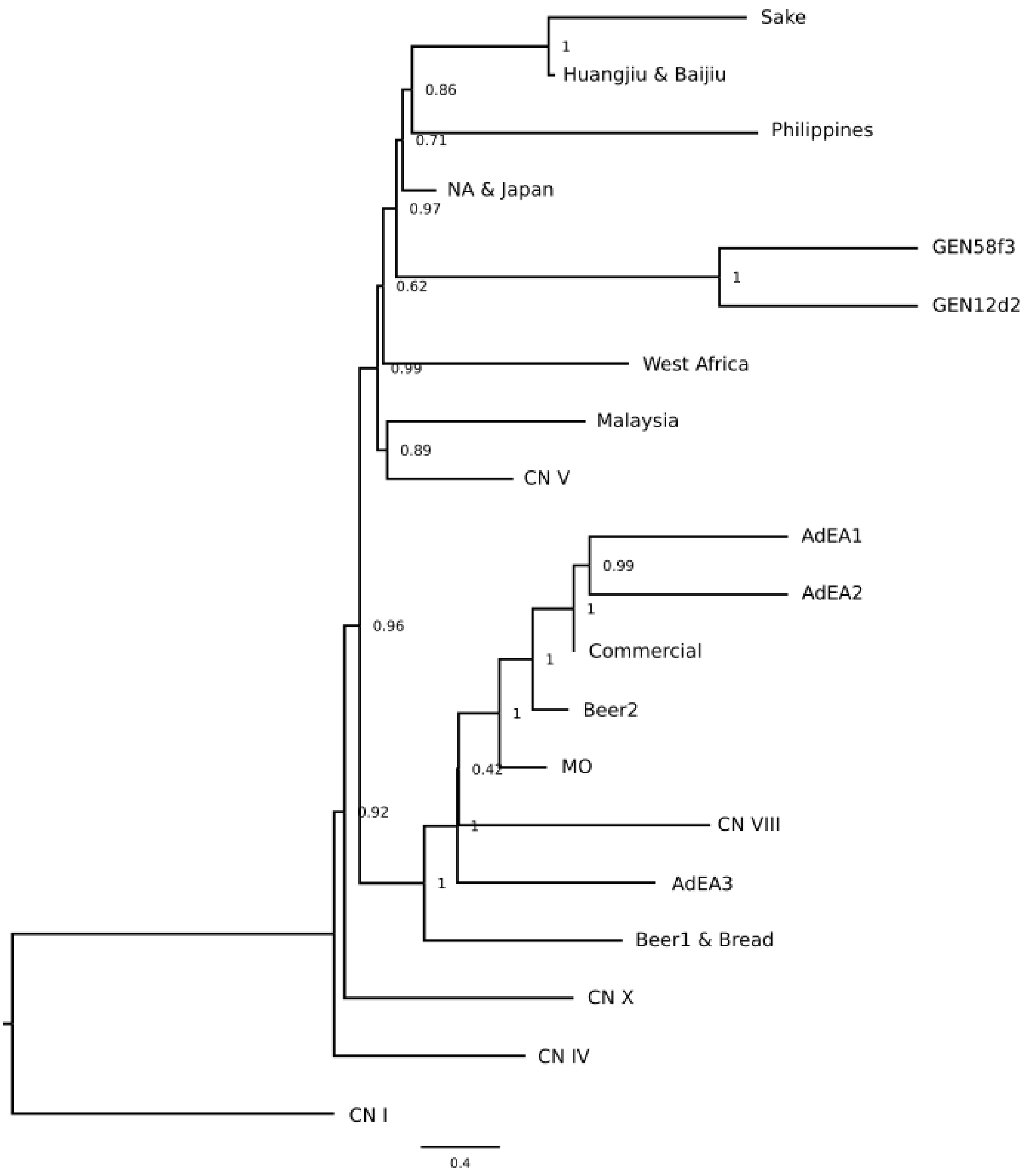


Sup Figure 16: Phylogenetic reconstruction from 691 single copy orthologs places isolates GEN12d2 and GEN58f3 as sharing a common ancestor with the NA & Japan clade. AdEA1, AdEA2 and AdEA3 are included as an indicator that although they contain admixture (~50% of the genome in AdEA3) they are still placed closer to domesticated lineages. Node posterior probability is shown. Scale is in coalescent units.

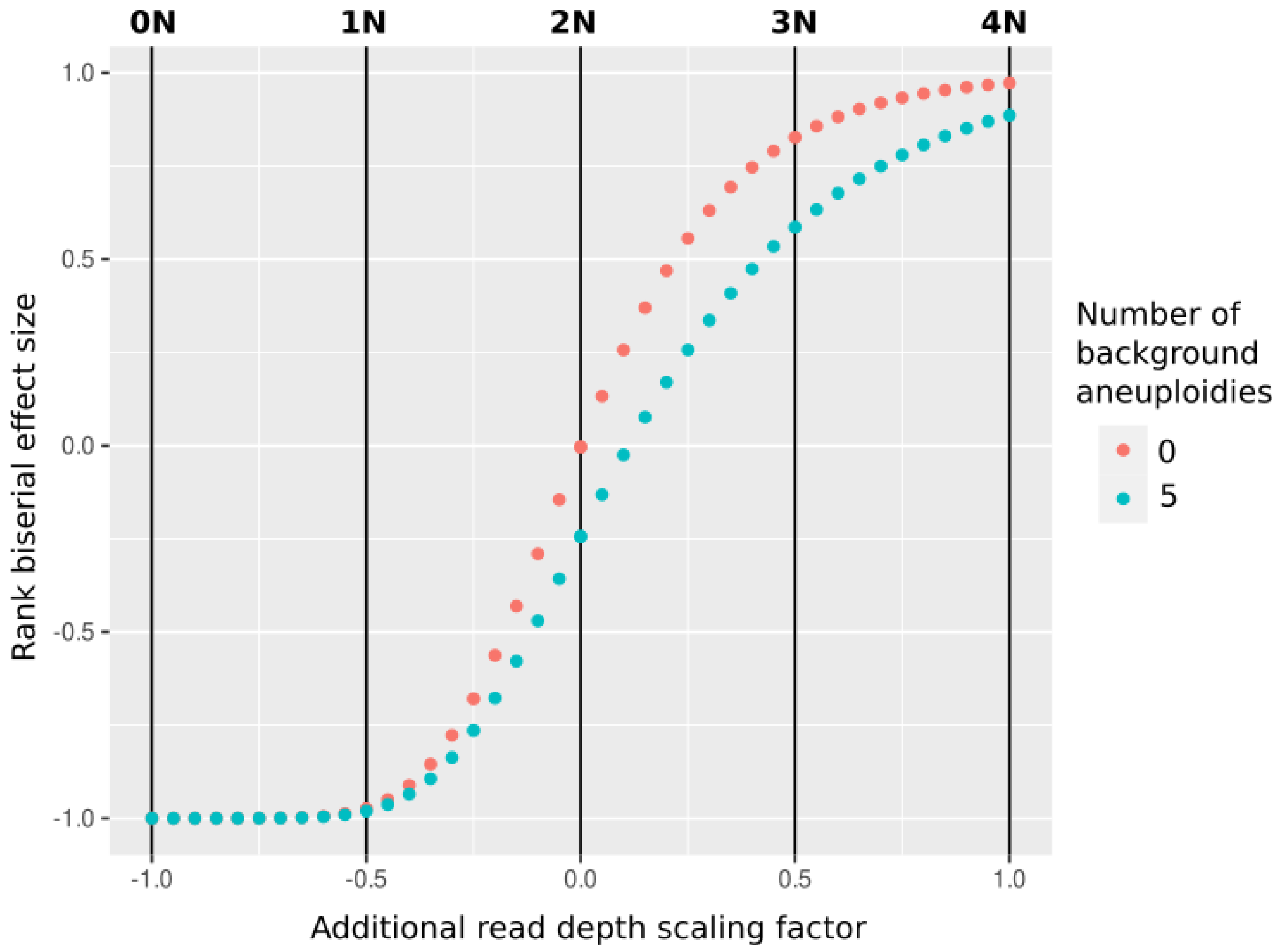


Sup Figure 17: Rank biserial effect size in simulated aneuploidies against genome wide read depth data. Aneuploidies were simulated in the genome wide background by scaling read depth for 0 and 5 chromosomes to 3N (ie, 1.5x chromosome read depth), aneuploid chromosomes were randomly selected from the empirically observed chromosome aneuploidy rate. Test chromosomes were scaled by a factor iterating through -1 to 1 (0N - 4N) by 0.1 to simulate read depth bias (2N read depth + 2N read depth x scaling factor).

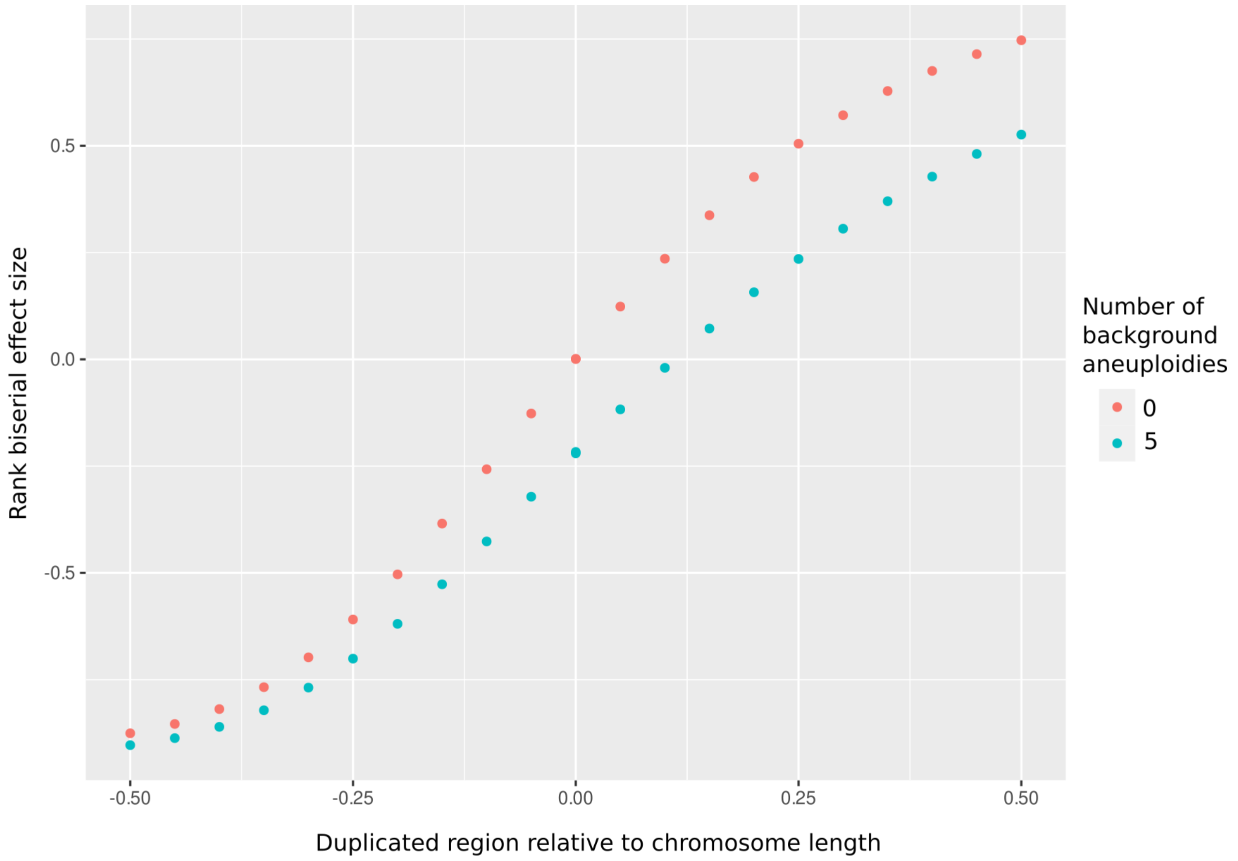


Sup Figure 18: Rank biserial effect size in simulated segmental duplications or transloctions against genome wide read depth data. Duplications were simulated in the genome wide background by scaling read depth for 0 and 5 chromosomes to 3N (ie, 1.5x chromosome read depth), aneuploid chromosomes were randomly selected from the empirically observed chromosome aneuploidy rate. Test chromosomes had a fraction of their length (-0.5 - 0.5) scaled by a factor of 1.5 (ie –0.5 has 1N for half the chromsome; 0.5 has 3N for half the chromosome).
